# Metastable Oscillatory Modes emerge from synchronization in the Brain Spacetime Connectome

**DOI:** 10.1101/2022.01.06.475196

**Authors:** Joana Cabral, Francesca Castaldo, Jakub Vohryzek, Vladimir Litvak, Christian Bick, Renaud Lambiotte, Karl Friston, Morten L. Kringelbach, Gustavo Deco

## Abstract

A rich repertoire of oscillatory signals is detected from human brains with electro- and magnetoencephalography (EEG/MEG). However, the principles underwriting coherent oscillations and their link with neural activity remain under debate. Here, we revisit the mechanistic hypothesis that transient brain rhythms are a signature of metastable synchronization, occurring at reduced collective frequencies due to delays between brain areas. We consider a system of damped oscillators in the presence of background noise – approximating the short-lived gamma-frequency oscillations generated within neuronal circuits – coupled according to the diffusion weighted tractography between brain areas. Varying the global coupling strength and conduction speed, we identify a critical regime where spatially and spectrally resolved metastable oscillatory modes (MOMs) emerge at sub-gamma frequencies, approximating the MEG power spectra from 89 healthy individuals at rest. Further, we demonstrate that the frequency, duration, and scale of MOMs – as well as the frequency-specific envelope functional connectivity – can be controlled by global parameters, while the connectome structure remains unchanged. Grounded in the physics of delay-coupled oscillators, these numerical analyses demonstrate how interactions between locally generated fast oscillations in the connectome spacetime structure can lead to the emergence of collective brain rhythms organized in space and time.

## Introduction

The human brain is one of the most complex networks in nature, exhibiting a rich repertoire of activity patterns organized not only in space and time but also in the frequency domain. Indeed, rhythmicity is a central property of brain function – and perhaps of all biotic self-organisation: from fast gamma activity in neurons to the life-cycle itself^1-4^. Within the broad range of oscillations emerging at frequencies between 0.05 Hz and 500 Hz, the oscillations detected extracranially with electro- and magnetoencephalography (EEG/MEG) in resting humans typically peak between 0.5 and 30 Hz, being categorized as delta (∼0.5-4 Hz), theta (∼4-8 Hz), alpha (∼8-13 Hz) and beta (∼13-30 Hz)^5^. Notably, these oscillations lock in phase over long distances, generating metastable spatial topographies lasting up to a few hundred milliseconds^6-8^.

Falling significantly below the range of frequencies generated in local neuronal networks by feedback inhibition (>35Hz, in the gamma frequency range), it is generally agreed that sub-gamma oscillatory activity does not have a purely local origin and is associated with synchronization between distant neural assemblies^9-14^. Notably, there is a relation between the distance over which synchronization is observed and the frequency of the synchronized oscillations^15-17^. Specific brain circuitries, including among others the thalamocortical loop, have been proposed to play a role in the generation of rhythmic activity^18-20^, which appears disrupted in neurological/neuropsychiatric disorders ^1,21^. Still, the fundamental mechanisms driving the spontaneous emergence of short-lived spatially and spectrally resolved oscillatory patterns remain unclear^9,22-24^.

Given the spatial distance and the finite propagation speed, interactions between brain areas are intrinsically time-delayed, which can manifest in network activity in the frequency domain. Indeed, delay-coupled limit-cycle oscillators have been demonstrated to synchronize at frequencies slower than the natural frequency of the oscillators, leading to a form of *collective* frequency emerging from synchronization mechanisms^25,26^. Briefly, when *N* phase oscillators – with natural frequency *ω* – are coupled together with a time delay *τ*, they synchronize at a delay- and interaction-dependent collective frequency *Ω* given by *Ω* =*ω* /(1+*K*\**N*\**τ*), where *K* is the global coupling strength^25^. However, this phenomenon has so far only been demonstrated for networks of limit-cycle oscillators^25^, and it is unclear how it generalizes to systems where oscillations are not self-sustained, but instead emerge only transiently.

Computational models have proved helpful for demonstrating how the brain’s complex network structure of long axonal projections connecting brain areas – the so-called structural connectome^27^ - can *shape* brain activity in space and time^28-36^. Particularly, simulations of oscillatory units interacting in the connectome reveal a critical regime where different subsets of units temporarily synchronize and desynchronize, leading to transiently correlated activity across spatially segregated units^30,37-39^. This reinforces the hypothesis that long-range functional connectivity between brain areas is driven by synchronization mechanisms^24,40-44^. Importantly, when considering realistic time delays in the Kuramoto model of coupled phase oscillators, periods of increased synchrony are accompanied by increased power at slower frequencies, generating spatially-organized band-limited power fluctuations similar to the ones captured with MEG^38^. While these numerical results revealed the critical role of time delays to generate collective oscillations at reduced frequencies, it remains to be verified whether this phenomenon holds in the more realistic setting, wherein local oscillations have fluctuating amplitude – which is neglected in the Kuramoto model –, as observed empirically in electrophysiological recordings of neural activity^45,46^. Furthermore, understanding the parameters that control the duration, size and occupancy of collective oscillations is crucial to inform the prediction of therapeutic strategies aimed at modulating dysfunctional oscillatory brain activity.

To address these fundamental questions, we build a phenomenological brain network model with realistic connectivity and time delays, where each node is described by a Stuart-Landau oscillator operating in the subcritical regime, i.e., responding to a stimulus with an oscillation with decaying amplitude ^34,35,47^. As the amplitude dynamics introduces an additional degree of complexity, it needs to be verified if the analytic predictions made for coupled limit-cycle oscillators^48^ (valid for phase oscillators or supercritical Stuart-Landau oscillators) still hold^49^. Selecting 40Hz as a typical frequency of gamma oscillations, we set all units with identical natural frequency to exclude additional effects of frequency dispersion^50,51^, and perturb all units with uncorrelated white noise, considering that units resonate at their natural frequency in the presence of background noisy activity^52^. Assuming the generalizability of collective synchronization frequencies to delay-coupled damped oscillators, we hypothesize to identify a critical range of global model parameters (global coupling and conduction speed) where metastable synchronization generates the transient emergence of sub-gamma collective oscillations, approximating features of human MEG recordings.

## Results

### Dynamical regimes of the brain network model

The reduced brain network model comprises *N*=90 dynamical units representing anatomically defined brain areas coupled according to a normative structural connectome of the human brain (see *Methods – Structural Connectome*) with reciprocal (i.e., bidirectional/symmetric) coupling *C*_*N×N*_ and distance *D*_*N×N*_ matrices (Figure 1a). Each unit is described by a Stuart-Landau oscillator operating in the subcritical (underdamped) regime, such that when perturbed it decays to a fixed-point equilibrium with a damped oscillation at a natural frequency *ω* (Figure 1b), in contrast with the supercritical regime, where the oscillations are in a limit cycle (Figure 1c, see *Methods* and *Supplementary Note 1*).

**Figure 1.**
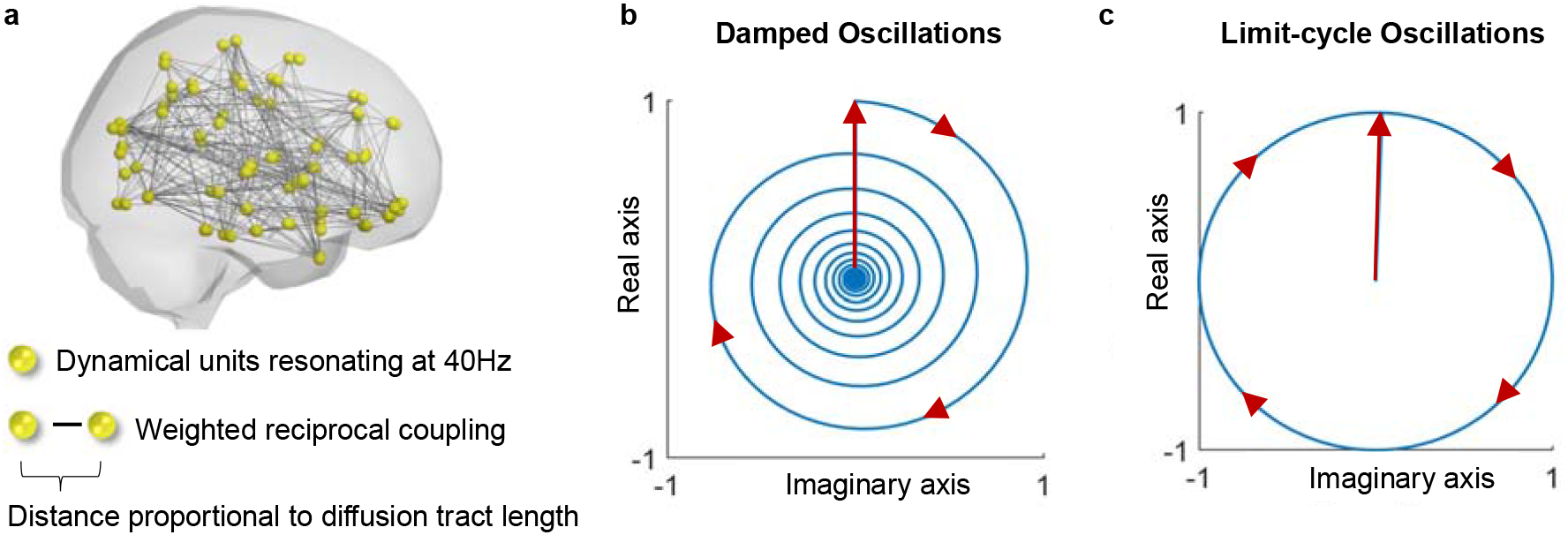
**a –** The phenomenological brain network model consists in N=90 nodes representing brain regions with links representing diffusion tracts between them. **b** – A Stuart-Landau oscillator in the subcritical regime responding to perturbation (vertical arrow) with an oscillation with decaying amplitude. **c** – In the supercritical regime, the Stuart-Landau oscillator enters a limit-cycle (with constant amplitude), approximating a phase oscillator.

To verify that novel frequencies emerge purely from delayed interactions, the natural frequency of all units is set at *ω* = 40Hz (representing the resonant frequency of isolated neural masses driven by feedback inhibition) and each unit is perturbed with uncorrelated white noise. The model – represented mathematically by a system of stochastic delay coupled differential equations – is solved numerically for two parameter ranges: the global coupling strength, *K*, which scales all pairwise connections, and the mean conduction delay, ⟨*τ*⟩, which scales the time delays between units in proportion to the diffusion tract lengths (Figure 2, see Methods for details).

**Figure 2.**
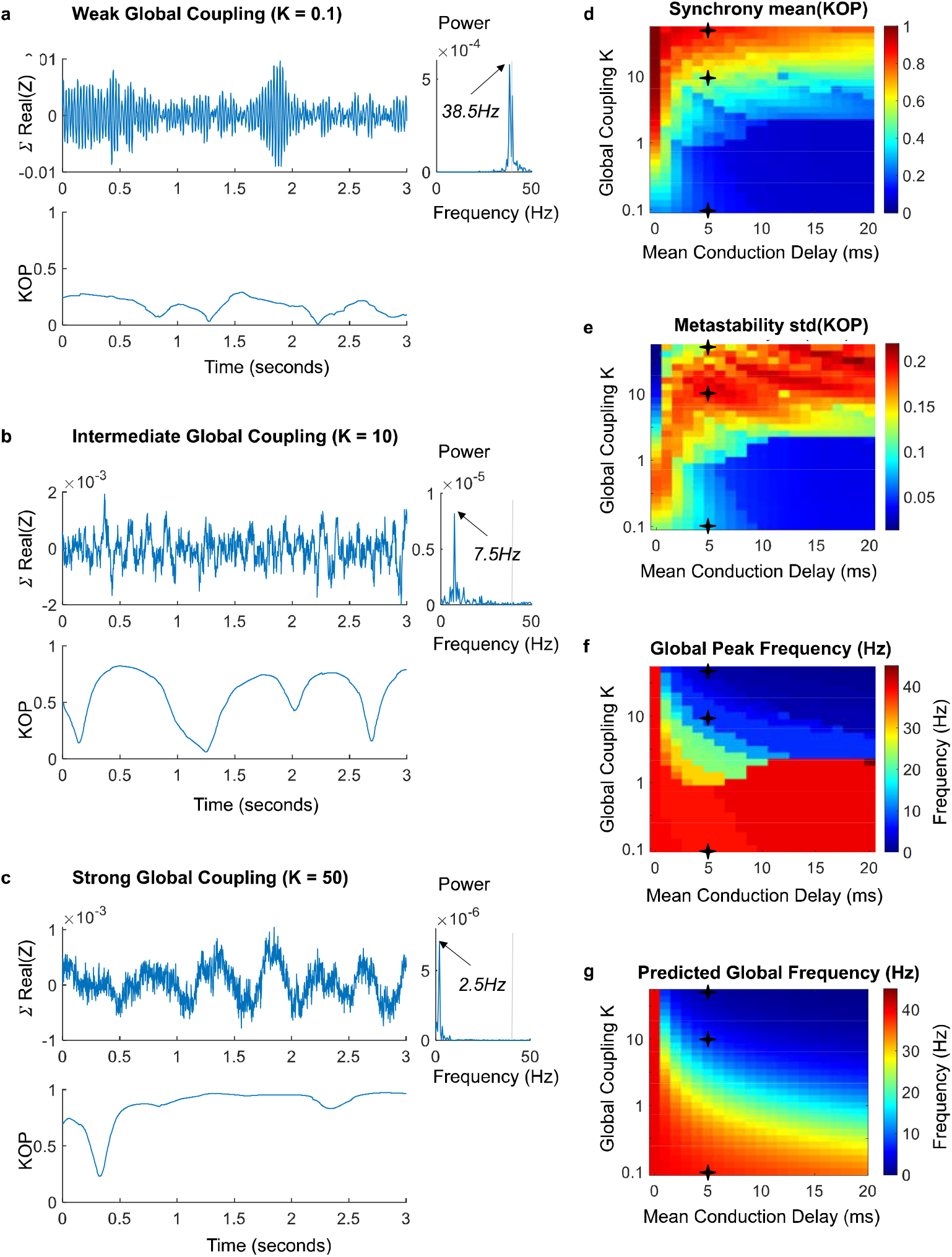
Collective oscillations emerge at reduced frequencies from time-delayed synchronization. The system of N=90 coupled oscillators, Z, was simulated for 50 seconds in the presence of white noise, varying only two global parameters: the Global Coupling K (increasing exponentially to better capture the effect of delays) and the conduction speed, which scales the Mean Conduction Delay. **a**,**b**,**c** – To illustrate the effect of the coupling strength in the frequency of synchronization, the collective signal given by 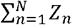. with N=90 is reported for three levels of global coupling, keeping the same mean conduction delay of 5 milliseconds. The corresponding power spectra are reported on the right of each plot, and the Kuramoto Order Parameter (KOP) is reported below. For weak coupling (**a**) the simulated signal exhibits oscillations peaking close to the node’s natural frequency. For intermediate coupling (**b**), weakly stable synchronization generates transient oscillations at reduced frequencies. For strong coupling (**c**), global synchronization becomes more stable, and all units are entrained in a collective oscillation at a reduced frequency. For intermediate coupling, fluctuations in the order parameter are indicative of metastability. **d-g** – For each simulation across the parameters explored, we report: (**d**) the mean of the KOP (referred to as Synchrony); (**e**) the standard deviation of the KOP (referred to as Metastability^54^); (**f**) the peak frequency of the simulated collective signal; (**g**) the synchronization frequency predicted analytically, showing agreement with simulation results for sufficient synchrony.

The synchrony degree of the system, evaluated using the Kuramoto Order Parameter (KOP), is modulated by the global coupling strength *K*: for weak coupling, the synchrony is low, and all units exhibit oscillations close to the natural frequency *ω* (Figure 2a). In the critical range between incoherence and full synchrony, periods of weakly stable synchronization drive slow fluctuations in the KOP (Figure 2b). For sufficiently strong coupling, all units tend to synchronize at a global collective frequency *Ω*, which, in the presence of time-delays, is distinct from the natural frequency *ω* (Figure 2c).

Observing the levels of synchrony and metastability across the range of parameters explored (Figure 2d-e), we find that the critical value of K above which the system can synchronize increases logarithmically with the mean delay, in line with analytic predictions for coupled oscillators with heterogeneous delays^53^ (see Supplementary Note 2 and Supplementary Figure 5). When synchronization occurs in the presence of delays, we observe a sharp decrease in the global peak frequency (Figure 2f), closely approximating the analytic prediction given by *Ω* =*ω* /(1+*K*\**N*\*⟨*τ*⟩) (Figure 2g, see also Supplementary Note 2 and Supplementary Figure 6).

These findings serve to verify that the phenomenon of synchronization at reduced collective frequencies is not restricted to coupled phase oscillators and generalizes to units in the subcritical regime, where damped oscillations emerge in response to perturbation (Supplementary Figure 7). Further, it demonstrates that the peak frequency of synchronization can be predicted analytically from global variables such as the mean natural frequency *ω*, the number of units *N*, the coupling strength *K*, and the mean delay ⟨*τ*⟩. The robustness of this prediction to distributed natural frequencies is reported in Supplementary Figure 8.

### Simulations reveal spectral features of human brain activity

One characteristic feature of MEG (and EEG) signals from healthy humans at rest is the transient emergence of oscillations in the alpha frequency range (∼8-13Hz), resulting in a peak in the power spectrum whose prominence varies strongly across people (see Figure 3a for the normalized power spectrum of MEG signals from 89 healthy young adults resting with eyes open from the Human connectome Project open-source database; details in *Methods* section, individual power spectra reported in Supplementary Figure 9).

**Figure 3.**
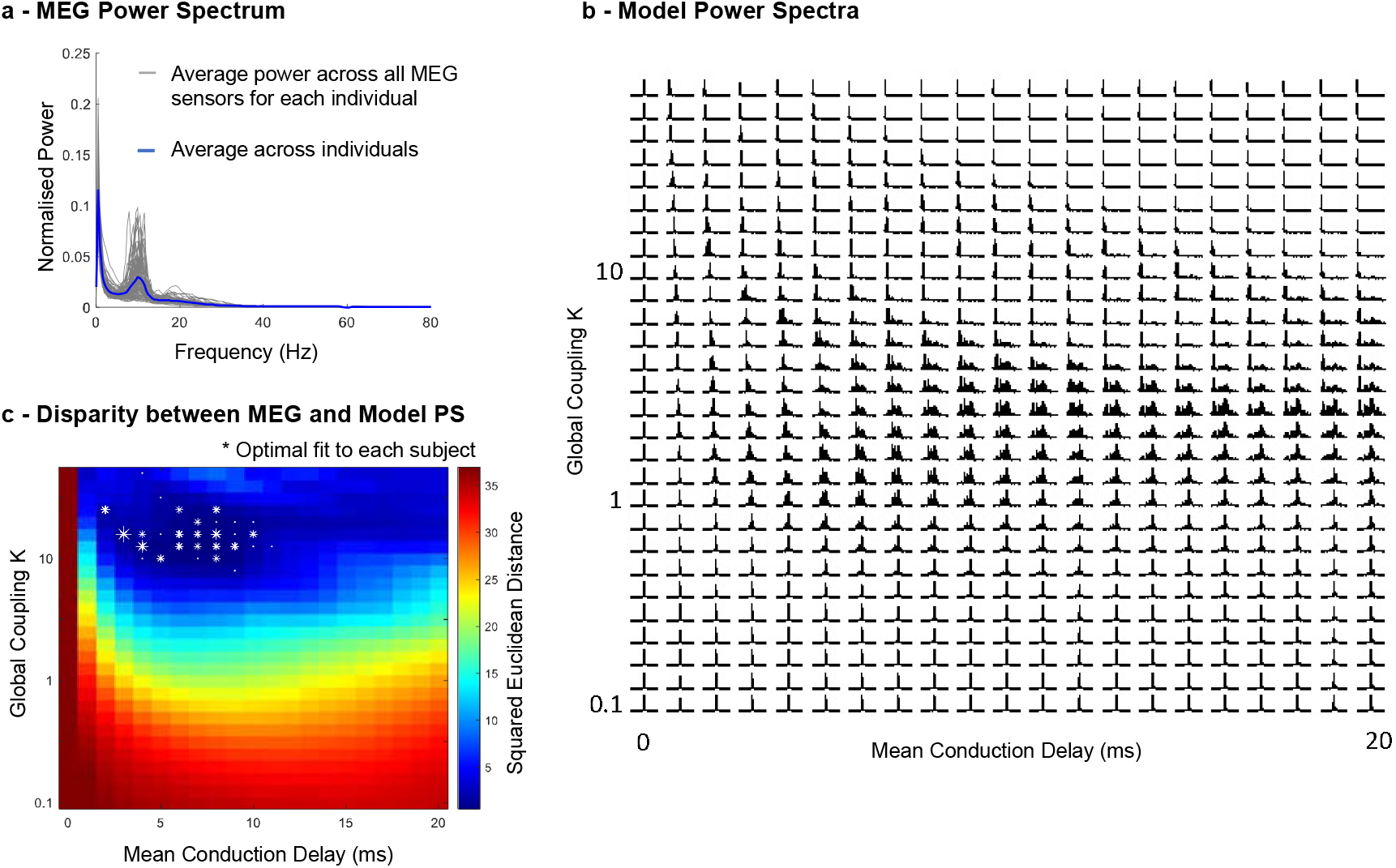
Approximation of human magnetoencephalography (MEG) power spectra (PS) in a critical range of parameters. **a** - MEG power spectra from 89 healthy young adults resting with eyes open from the open-source database of the Human connectome Project. The average power spectrum across individuals is reported in blue. **b** – For each pair of parameters, the power spectra of the simulated signals (averaged across units and normalized between 0 and 80Hz) is reported. **c** – Squared Euclidean distance between the MEG power spectrum averaged across all sensors and subjects and the power spectrum of the simulated signals. Asterisks indicate the sets of parameters that optimally approximate the MEG power spectra of each of the 89 individuals (size scaled according to the number of subjects in each point).

We find that the brain network model approximates the average MEG power spectrum of awake resting subjects within the critical region of high metastability where synchronization occurs at reduced collective frequencies (comparing Figure 3c with Figure 2e-f). In detail, for each pair of model parameters we calculate the squared Euclidean distance between the power spectrum of the simulated signals (Figure 3b) and the MEG power spectrum averaged across all sensors and all subjects (Figure 3a), revealing the greatest disparity when no delays are considered or if the global coupling is too weak (see Methods for details).

Given the observed (and well-established) variability between MEG power spectra across individuals (Figure 3a), we investigate the extent to which this variability can be associated with changes in global model parameters, while keeping the structural connectivity unchanged. To do so, we identify the pair of model parameters that approximates the individual MEG power spectra of each of the 89 participants, falling in 29 pairs of parameters (white asterisks in Figure 3c, see also Supplementary Figure 10). Notably, this reveals a confined region in parameter space for a range of average delays ⟨*τ*⟩ of 2 to 11 milliseconds, with slight changes in the coupling strength and conduction speed maximizing the fit to individual MEG power spectra, while the structural connectivity remains unchanged. These results do not exclude the role of individual variability in structural connectivity across subjects but reveal additional parameters that modulate a network’s frequency spectrum. This serves to demonstrate that the same connectome structure can support distinct activity patterns depending on global model parameters, with longer/shorter time delays and stronger/weaker coupling inducing shifts in the peak frequency and modulating the distribution of power across the spectrum (Figure 3b).

### Metastable Oscillatory Modes emerge from weakly stable cluster synchronization

In the range of parameters where the model optimally approximates the power spectrum of MEG signals, fluctuations in the magnitude of the order parameter are driven by metastable cluster synchronization. In other words, when the coupling is strong, but not sufficiently strong to stabilize full synchronization, some subsets of units that are more strongly connected together (i.e., clusters/communities) can engage in partially-synchronized modes that remain stable for a short period in time. Given the presence of time delays, these clusters do not synchronize at the natural frequency of the individual units (*ω* =40Hz), but instead synchronize at slower cluster-specific collective frequencies, leading to the emergence of metastable oscillatory modes (MOMs) at sub-gamma frequencies.

To detect the occurrence of MOMs and characterize them in space and time, we band-pass filter the simulated signals in 4 frequency bands (delta 0.5-4Hz, theta 4-8Hz, alpha 8-13Hz and beta 13-30Hz). In Figure 4, a coloured shade is added when the amplitude in each frequency band exceeds 5 standard deviations of the amplitude in that same frequency range detected in simulations without delays (see Supplementary Note 3, Supplementary Figure 11).

**Figure 4.**
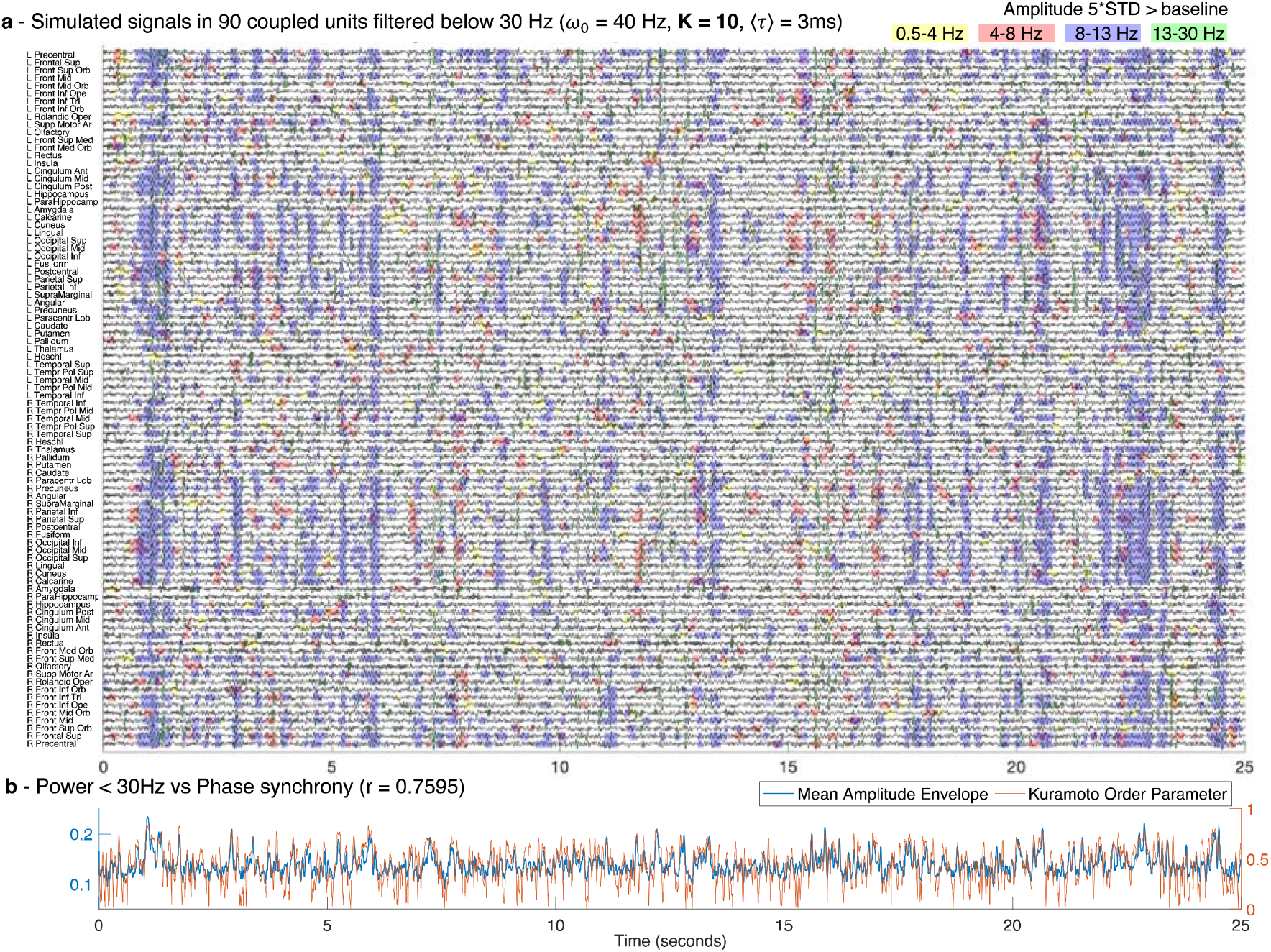
Sub-gamma oscillations emerge from weakly stable cluster synchronization. **a** – An example of the simulated signals in all 90 units plotted over 25 seconds, each representing a brain area from a brain parcellation template, filtered below 30 Hz to highlight the sub-gamma oscillatory activity typically detected with magnetoencephalography (MEG). Shades indicate the time points of increased power in the delta (yellow), theta (red), alpha (blue) and beta (green) frequency bands. For each frequency band, the threshold was defined as 5 standard deviations (STD) of the amplitude – in the same frequency bands – when no delays were considered. For the simulations, the resonant frequency, *ω*_0_, of all units was set to 40Hz, the conduction speed was tuned such that the average delay between units, ⟨*τ*⟩, was 3 milliseconds (ms) and the global coupling strength was set to K=10. **b** –The mean amplitude envelope (blue) of the filtered signals shown in (a) correlates with a Pearson’s correlation coefficient r=0.7595 with the phase synchronization evaluated by the Kuramoto Order Parameter (orange, right y-axis).

As shown in Figure 4, we find that MOMs are structured both in space and in time. Specifically, the units synchronizing together exhibit the simultaneous emergence of an oscillation at the same collective frequency, leading to the vertical alignment of shaded areas, particularly visible for the alpha frequency range in Figure 4a. Notably, for different sets of parameters, the configuration of Figure 4a changes strongly. Indeed, while for very weak coupling, almost no supra-threshold oscillations are detected (Supplementary Figure 12), for stronger coupling globally synchronized supra-threshold oscillations emerge transiently in the delta band (Figure 5a). For longer delays, oscillations are detected with a less definitive temporal alignment between brain areas (Supplementary Figure 13).

**Figure 5.**
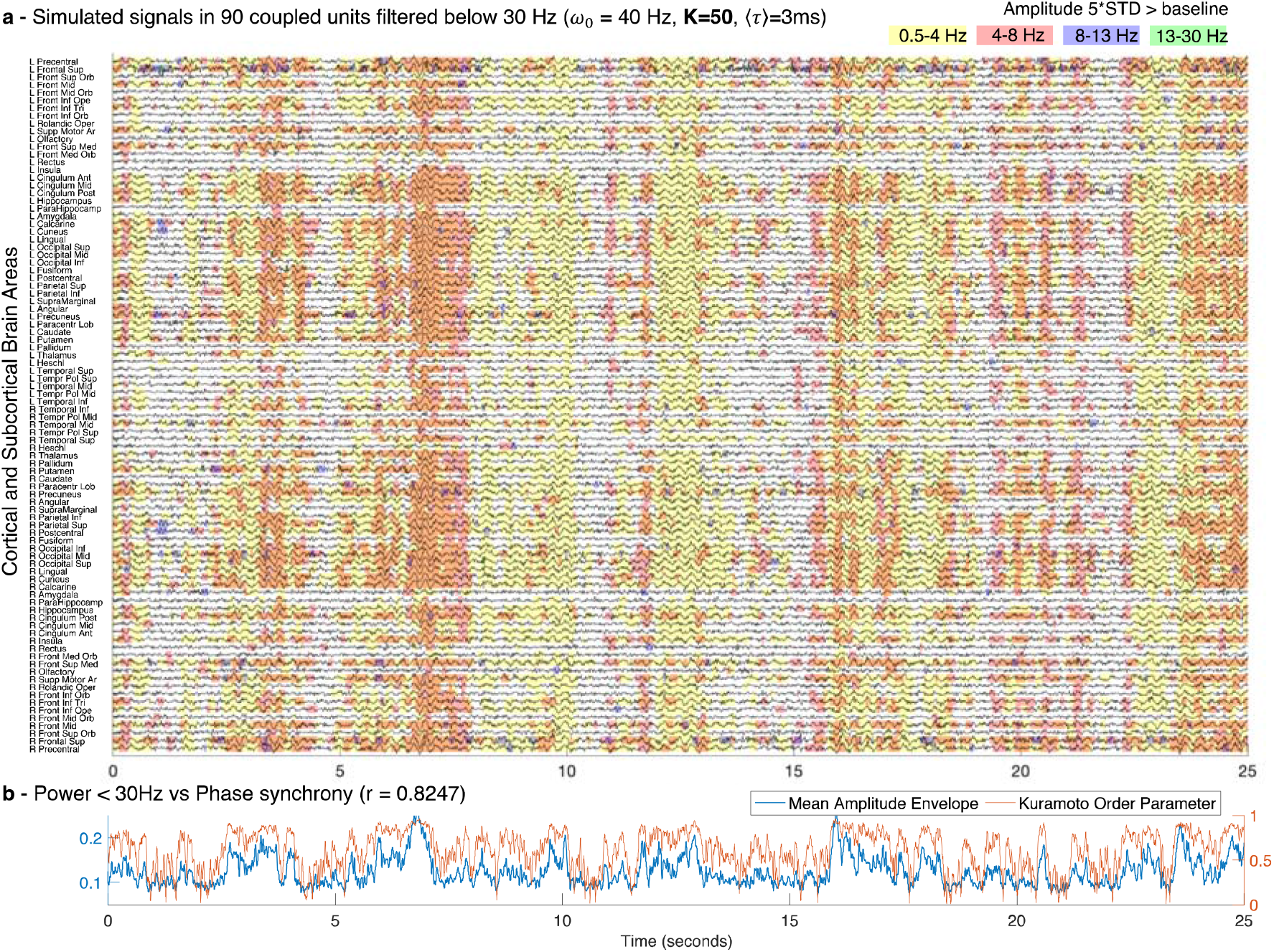
Global delta waves emerge for strong coupling. **a** – The simulated signals in all 90 units plotted over 25 seconds, each representing a brain area from a brain parcellation template, filtered below 30 Hz to focus on the sub-gamma oscillatory activity typically detected with magnetoencephalography (MEG). Shades highlight the time points of increased power in the delta (yellow), theta (red), alpha (blue) and beta (green) frequency bands. For each frequency band, the threshold was defined as 5 standard deviations (STD) from the amplitude in the same frequency bands when no delays were considered. These simulations were performed setting the resonant frequency of all units *ω*_0_ = 40Hz, the average delay between units, ⟨*τ*⟩ = 3 milliseconds (ms) and the global coupling strength was increased to K=50 with respect to the simulations shown in Figure 4. **b** –The mean amplitude envelope (blue) of the filtered signals shown in panel A correlates with r=0.8247 with the phase synchronization evaluated by the Kuramoto Order parameter (orange, right y-axis).

Furthermore, the power at sub-gamma frequencies is found to correlate strongly with the instantaneous phase synchronization evaluated by the KOP over time (r=0.7595 and r=0.8247 for Figure 4b and Figure 5b correspondingly). This demonstrates that the emergence of oscillations at sub-gamma frequencies in the simulations is modulated by fluctuations in the synchrony degree.

We further define quantitative metrics to characterize the MOMs emerging at different frequency bands for different sets of model parameters in terms of their duration (i.e., consecutive time that the power remains above threshold), their size (i.e., the number of units simultaneous displaying power above threshold) and occupancy (i.e., the proportion of time that the power is detected above threshold).

As can be seen in Figure 6, in the range of parameters where optimal fits to MEG data are obtained (Optimal Range), the alpha MOMs last longer, recruit more units and occur more often. Importantly, we demonstrate that global parameters, such as the coupling strength and the conduction speed, modulate the spatiotemporospectral properties of the whole system in a non-trivial way, while the dynamics at the local level and the underlying structural network remain unchanged.

**Figure 6.**
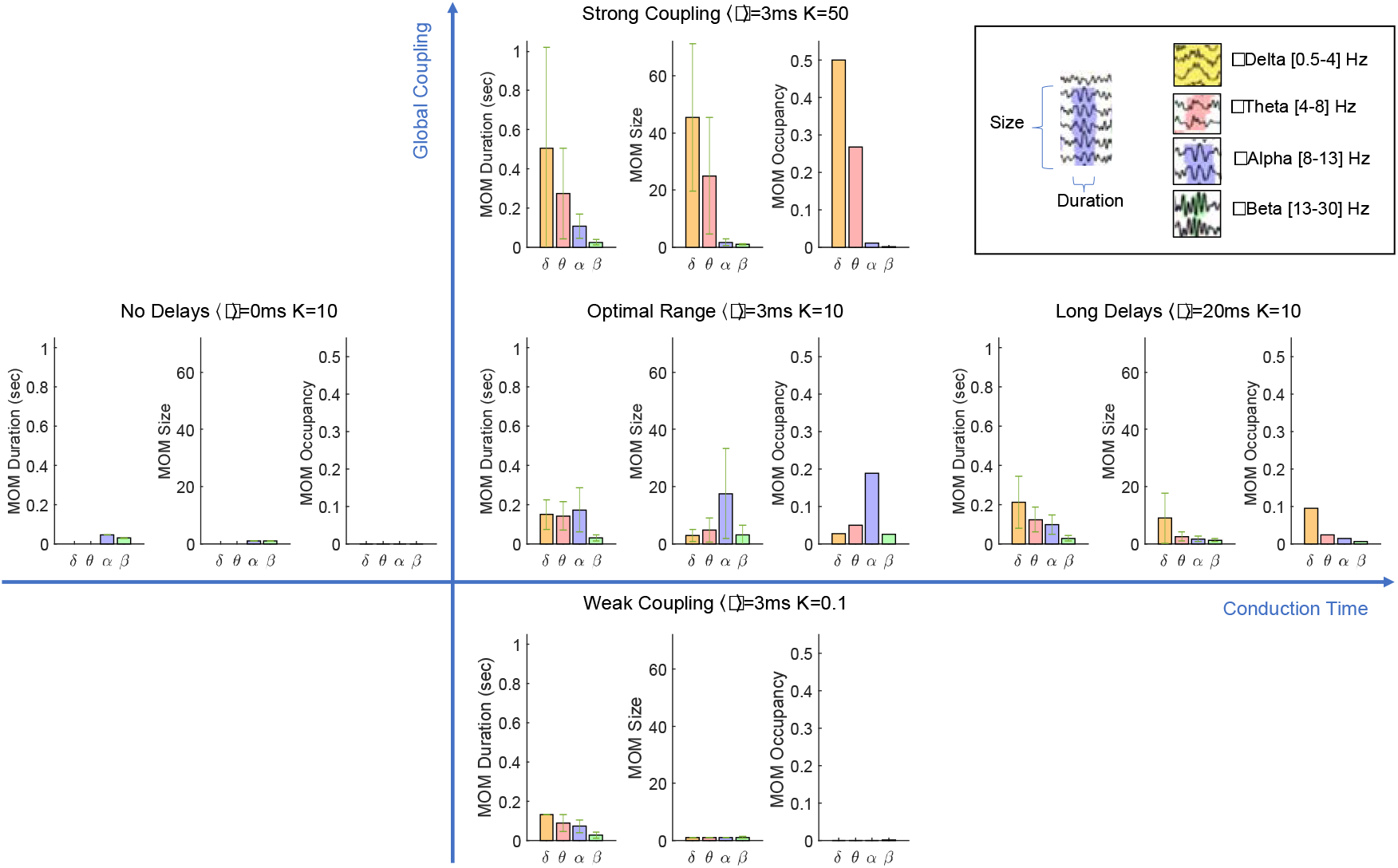
Characterization of metastable oscillatory modes (MOMs) emerging from the system. For different Global Coupling strength (K) and Conduction Delays ⟨τ⟩, MOMs are characterized in terms of duration (i.e., consecutive time that the power remains above threshold), size (i.e., the number of units simultaneous displaying power above threshold) and occupancy (i.e., the proportion of time that the power is detected above threshold over the entire simulation), for each frequency band. This demonstrates that the same network structure, i.e., the connectome, can exhibit different oscillatory modes organized in space and in time, depending on global parameters of the system. In the critical range of parameters (Optimal Range), oscillations in the alpha frequency band emerge more frequently and involve more units. Globally synchronized delta oscillations – as typically observed in states of reduced consciousness – are associated to an increase in the global coupling strength (Strong Coupling). Error bars represent 1 standard deviation. See also Supplementary Movie 1.

The implicit sensitivity to global model parameters is illustrated in Figure 7, where the emergence of supra-threshold oscillations in different frequency bands is represented in the brain at a single time point for five distinct sets of parameters. The evolution over time is shown in Supplementary Movie 1.

**Figure 7.**
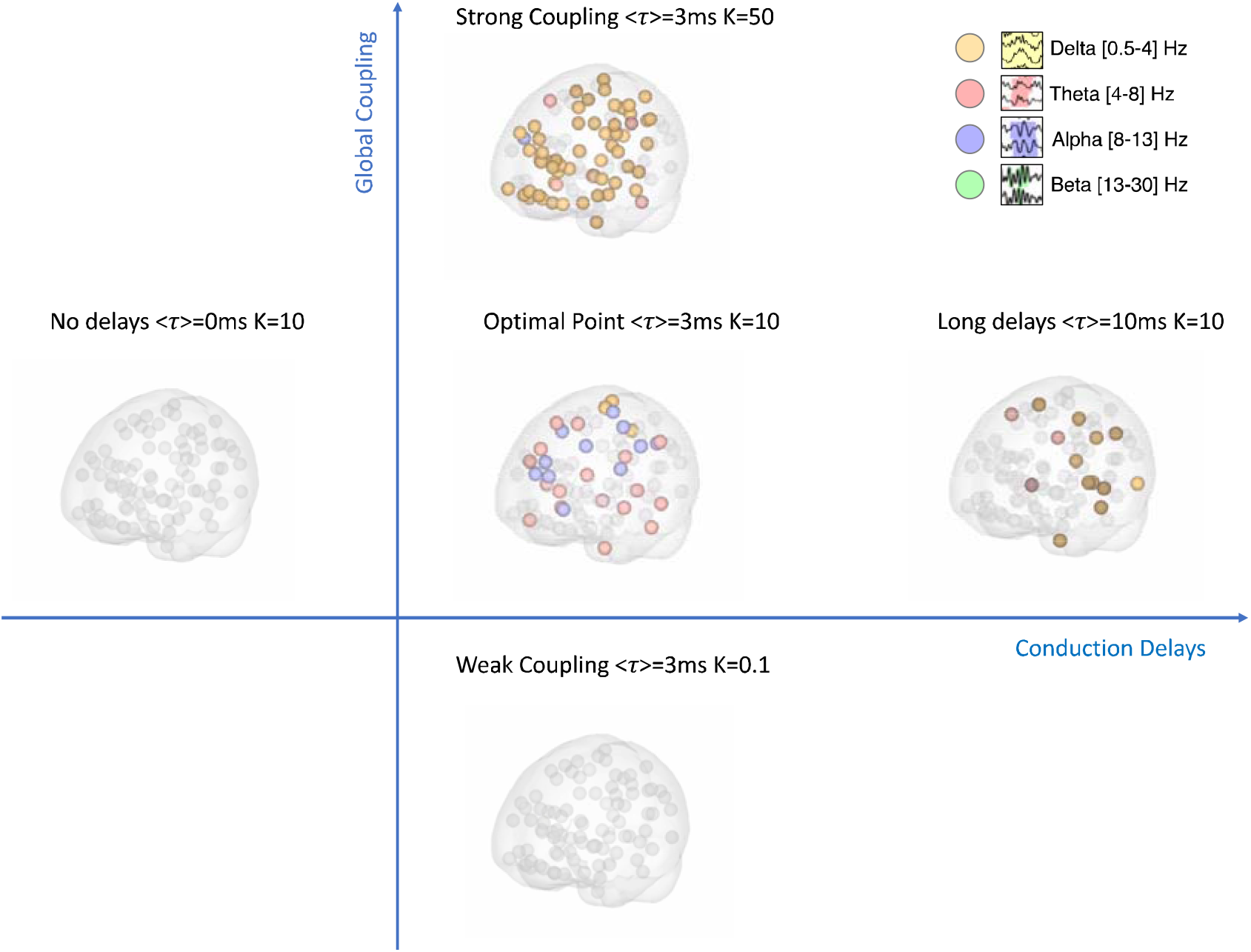
Metastable Oscillatory Modes (MOMs) emerge transiently from interactions in the Connectome spacetime structure only for sufficient coupling and conduction times. Each brain area is represented as a sphere located at its centre of gravity. A colour code is used to highlight the brain areas with power exceeding 5 standard deviations from the baseline power at a given time point. This image is a still frame from Supplementary Movie 1. While the structural connectome is the same for all simulations, MOMs only emerge at reduced frequencies in the presence of Conduction Delays (r) and for sufficient Coupling strength (K).

### Frequency-specific functional connectivity

To link with studies of functional connectivity in MEG, we further investigate how the model parameters modulate the correlation between the amplitude envelopes across frequency bands. To do so, we band-pass filter the signals in each frequency band, extract the amplitude of the Hilbert transform and report the envelope correlation matrices in Figure 8 for each frequency band and for four representative sets of model parameters. For weak coupling, the envelope correlations are close to zero (Pearson’s correlation coefficient cc<0.1 for all pairs of brain areas), indicating that the coupling is insufficient to drive functional connections between brain areas. For global parameters in the optimal range (here *K*=10 and ⟨*τ*⟩ =3ms), different brain areas exhibit correlated envelopes, with stronger correlations (up to cc=0.78) being detected in the alpha frequency range. In contrast, for strong coupling the functional connectivity in the alpha band is reduced (maximum pairwise correlation of cc=0.25), while the envelopes of delta and theta oscillations are strongly correlated across the brain (up to cc=0.89). Keeping the optimal range of global coupling, *K*=10, but increasing the delays to an average of ⟨*τ*⟩=20ms, envelope functional connectivity is detected mostly in the delta frequency range. This illustrates how, given the same underlying spacetime network structure (i.e., the matrices of coupling weights *C* and distances *D*), changes in global parameters strongly affect the envelope functional connectivity patterns at different frequency bands.

**Figure 8.**
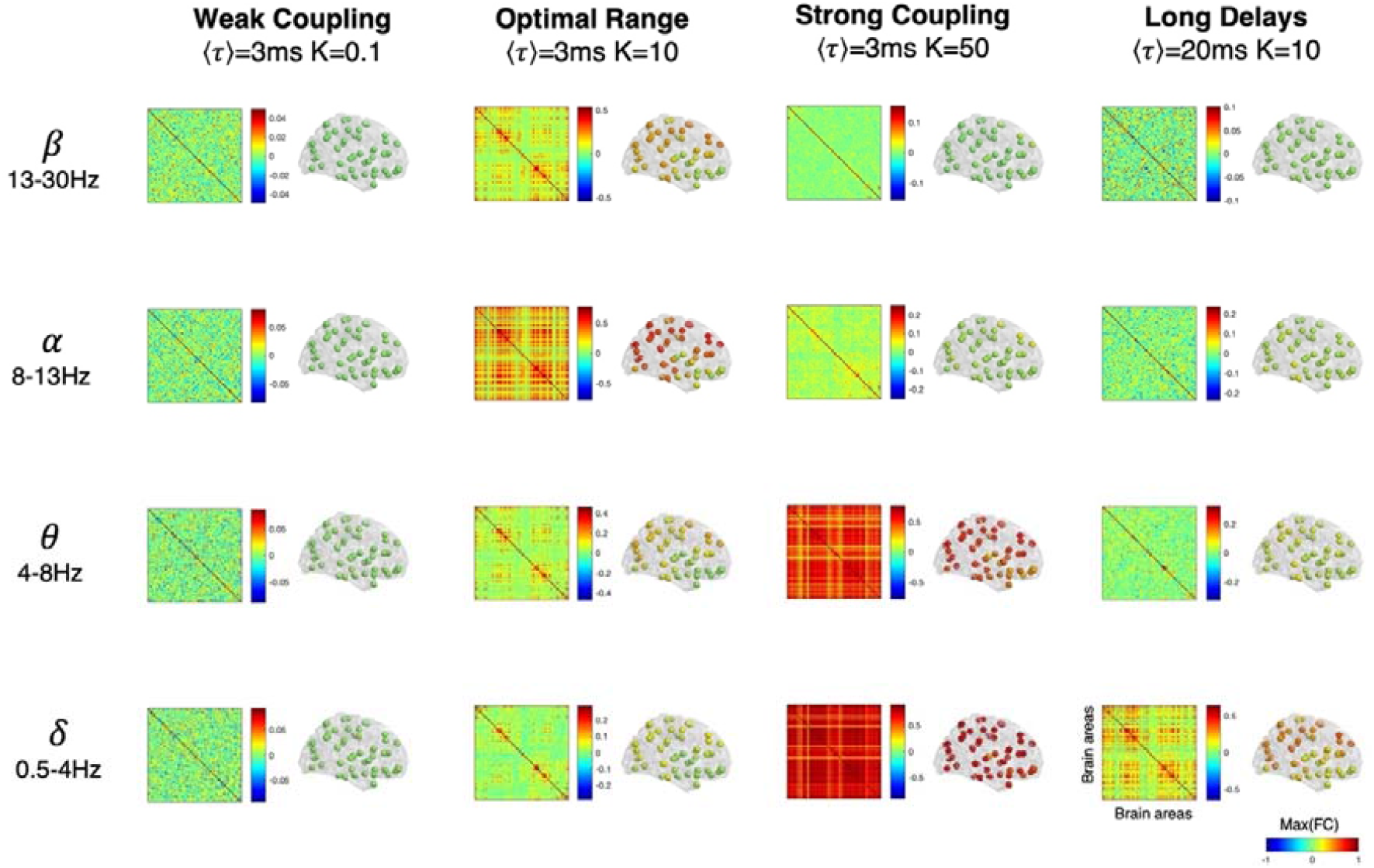
Influence of global model parameters in frequency-specific envelope functional connectivity patterns. For 4 simulations obtained with different Global Coupling strength (K) and Conduction Delays ⟨*τ*⟩, we report the frequency-specific functional connectivity (FC) estimated as the correlation matrices of the envelopes of signals band-pass filtered in the delta (δ), theta (θ), alpha (α) and beta (β) frequency bands. The colormap limits of the matrices are scaled by the maximum absolute correlation and centred at zero. Next to each matrix, each of the N=90 brain areas is represented as a sphere placed in its centre of gravity and coloured according to the maximum envelope FC to any other brain area (same colorbar applied to all spheres, scaled between −1 and 1).

To illustrate the level of functional connectivity across the brain, next to each correlation matrix in Figure 8, we represent each area as a sphere placed at its centre of gravity and coloured according to the strongest correlation with any other brain area. This shows that, for the optimal range of parameters, the areas exhibiting the strongest functional connectivity in the alpha band are distributed mostly in posterior and dorsal cortical areas, aligning with empirical observations of stronger functional connectivity in the alpha band in the visual and somatomotor systems. However, it is important to consider that the specific spatial configuration of functional connections is inherently dependent on the resolution and topology of the structural connectome, which is known to depend on the parcellation scheme and on the brain parts (i.e., cortical, subcortical) considered. In Supplementary Methods 1, we perform the same analysis on data simulated using a structural connectome including 200 cortical-only brain areas^55^. These results show that, while the phenomenology of MOMs is robust to changes in the parcellation scheme, the spatial specificity across frequency bands is sensitive to the parcellation scheme considered (Supplementary Figures 16 and 17). Most importantly, this analysis illustrates how frequency-specific functional connectivity patterns depend sensitively on global variables modulating the distributed dynamics, while the structural connectivity remains unchanged.

## Discussion

This work addresses the physical mechanisms underlying brain rhythms detected empirically, employing a reductionist perspective to ground the inner complexity of encephalographic signals to universal theoretical principles^56,57^. Approaching the problem from a macroscopic perspective, we focus on the emergent properties of interacting dynamical units, where the collective ensemble engages in functionally relevant activity patterns that cannot be inferred from the isolated units alone^58-61^.

Specifically, we first demonstrate the generalizability of a synchronization mechanism described for networks of delay-coupled *limit-cycle* oscillators to networks of delay-coupled *damped* oscillators (i.e., in the subcritical range of a Hopf bifurcation). This is important for the neuroscience field, since empirical electrophysiological recordings show that local field oscillations in the gamma frequency band are not limit-cycle oscillations (as considered in previous models using the Kuramoto of coupled oscillators^20^), but instead emerge only transiently. Therefore, the substantial reduction of brain areas to phase oscillators in Cabral et al. (2014) has raised concerns on the generalizability of the proposed mechanism to more realistic settings, given the demonstrated importance of considering the amplitude dynamics on the connectivity between phases ^48,49,62,63^.

Subsequently, we extend on previous brain network modelling works by demonstrating that the synchronization frequency can be approximated analytically from global model parameters, namely the number of units, the mean coupling strength, the average time delay between units, and the mean natural frequency of the units. Regarding the latter, we show that, in the presence of delays, the system is less sensitive to the spread of frequencies across units, in line with theoretical predictions^25^ (Supplementary Figure 8).

These insights are crucial to explain the macroscopic spatiotemporally organized oscillatory signals detected with EEG/MEG at sub-gamma frequencies, without explicitly introducing these oscillations in the model^64^. Here, we consider that only gamma-frequency oscillations can be generated at the local neuronal level, with power at other frequencies resulting purely from synchronization with time delays. Furthermore, we demonstrate the impact of global model parameters in the modulation of frequency-specific collective oscillations emerging across space and time. The detailed characterization of metastable oscillatory modes in terms of number of units synchronizing together, duration and occupancy provides a new framework to analyse collective brain oscillations complementary to frequency-specific envelope functional connectivity analysis.

Our hypothesis is endorsed using a phenomenological brain network model, reduced to its key essential ingredients to allow efficient numerical approximations to analytic predictions, but at the same time sufficiently complex to allow a fair approximation of MEG spectral features. The deliberate reductionist perspective inherent in this brain network model is intended to link with theoretical works on delay-coupled oscillatory systems^25,53,65,66^. Towards this end, we consider identical units with same natural frequency, same damping coefficient and same noise level, coupled in the structural connectome. Therefore, we focus solely on the effects of global variables, namely the global coupling *K* and the mean conduction delay ⟨*τ*⟩in the emerging synchronization phenomena. To establish the construct validity of our numerical simulations, we show that the peak synchronization frequency can be approximated by the analytic prediction derived for synthetic networks of coupled Kuramoto oscillators with time delays ^25^. Further, in line with theoretical predictions^53,67^, we find that the complex spacetime topology of the structural connectome widely expands the critical border between incoherence and global synchrony where fluctuations in the order parameter are indicative of metastability^25^. Despite its simplicity, this model provides a robust framework to test a theoretically grounded mechanistic scenario for the spontaneous formation of frequency-specific long-range coherence in complex networks.

While the investigation of mechanistic principles and control parameters benefits from reduced complexity, adding heterogeneity is certainly needed to improve the fitting to real brain activity from individuals in different conditions. Building up on these fundamental aspects, additional degrees of complexity can be added to the model, namely by considering more fine-grained connectome structures, considering non-homogeneous intrinsic frequencies and damping properties, or even replacing the noisy input by dynamic concentration patterns to mimic local neuromodulatory effects. Further, given the potential generalizability of this synchronization mechanism, we expect our analysis may provide valuable insight to interpret some of the complex self-organizing phenomena emerging in more realistic biophysical models of neural networks^68,69^ for which a precise analytic prediction cannot be solved.

Our findings reinforce the idea that conduction delays – often neglected in network models of whole brain activity due to the added complexity – play a crucial role in shaping the frequency spectrum of coupled oscillatory systems. Although the frequency of the oscillations considered herein is relatively fast with respect to the ultra-slow fluctuations (<0.1 Hz) detected with functional Magnetic Resonance Imaging (fMRI), it is important to highlight that metastable synchronization drives power fluctuations on ultra-slow timescales, and therefore, even relatively short time delays can significantly modulate spontaneous activity at ultra-slow time-scales. We note that for the numerical integration of stochastic delay differential equations to be stable and align with analytic predictions, the time step for numerical integration needs to be sufficiently small and a running history needs to be saved for the length of the maximum delay between units, which significantly increases the computation times when compared to simulations where delays are neglected (here the numerical results were found to stabilize for dt *≤* 10^−4^ seconds, see Supplementary Figures 18-20).

The discovery of multistability in systems of delay-coupled oscillators, initially described in 1999 by Young and Strogatz^65^ and extended to heterogeneous delays in 2009 by Lee, Ott and Antonsen^53^, was crucial to develop the theoretical hypothesis behind this work, opening grounds to speculate that this phenomenon may be related to the maintenance of the right balance between integration and segregation in living brains^70,71^. Beyond the range where the model best approximates healthy awake brain activity, we find that higher coupling enhances global order, where the whole brain displays slow coherent oscillations in the delta-range (0.5-2Hz), which nicely approximate the most powerful brain rhythms detected during unconscious states such as slow-wave sleep, coma or anaesthesia. On the other hand, operating at weaker coupling hinders the formation of MOMs at sub-gamma frequencies, altering the spectral profile similarly to what is observed in M/EEG recordings of patients with neuropsychiatric disorders associated to disconnection, such as schizophrenia, where the power in alpha appears to be significantly reduced^72-75^. Such abnormal interactions within cortico-subcortical oscillatory networks may emerge from specific local deregulation or neural circuit disruption^76^. However, how a local change may alter the communication between brain-areas and brain network dynamics remains an open question. Overall, these results are aligned with recent works proposing that spontaneous transitions between multiple space-time patterns on complex networks provide a solid theoretical framework for the interpretation of the non-stationary but recurrent macroscopic patterns emerging spontaneously in brain activity, and ultimately supporting brain function^77,78^. From a technical perspective, it may be surprising that this kind of itinerant dynamics emerges under symmetrical coupling between nodes; in the sense that asymmetric coupling is normally required for breaking detailed balance – and engendering stochastic chaos of the sort described above. However, the dynamics of each node are generated with asymmetric Jacobians, suggesting that symmetry breaking of intrinsic connectivity is a sufficient condition for the nonequilibrium dynamics that characterise real brains.

While metastability appears to be crucial for brain function, the specific role of MOMs to support cognitive functions remains unclear^4,24,71,79,80^. One possibility is that the areas engaged in a MOM are directly involved in long-range functional integration, but another is that these areas are inhibited by entering in a collective low-energy mode^13,41^. Shedding some light on this open question, we find that synchronization with delays induces not only a shift to slower frequencies but also a decrease in amplitude, in line with theoretical studies reporting amplitude death in systems with distributed delays^81^ (see the vertical axes in Figure 2a-c, top panels). From a ‘metabolic’ perspective, this shows that MOMs can be approached as ‘low energy modes’ with respect to high power gamma oscillations, providing a physical explanation for the emergence of the so-called ‘idle rhythms’^82^. Although the functional implications of this mechanism are beyond the scope of this work, we expect it will provide fertile grounds for the formulation of novel falsifiable predictions to be further tested. Moreover, these findings give room to further investigations of how local perturbations can affect the spatiotemporospectral dynamics on the macroscopic scale, to gain insight on the mechanisms of action of perturbative strategies such as transcranial magnetic stimulation or deep brain stimulation.

## Methods

### Ethics statement

All human data used in this study is from the public repository of the Human Connectome Project (HCP)^83^ (https://www.humanconnectome.org), which is distributed in compliance with international ethical guidelines.

### Structural connectome

The *NxN* matrices of structural connectivity, C, and distances, D, used for the network model were derived from a probabilistic tractography-based normative connectome provided as part of the leadDBS toolbox (https://www.lead-dbs.org/)^84^. This normative connectome was generated from diffusion-weighted and T2-weighted Magnetic Resonance Imaging (MRI) from 32 healthy participants (mean age 31.5 years old ± 8.6, 14 females) from the HCP. The diffusion-weighted MRI data was recorded for 89 minutes on a specially-designed MRI scanner with more powerful gradients then conventional scanners. The dataset and the acquisition protocol details are available in the Image & Data Archive under the HCP project (https://ida.loni.usc.edu/). DSI Studio (http://dsi-studio.labsolver.org) was used to implement a generalised q-sampling imaging algorithm to the diffusion data. A white-matter mask, derived from the segmentation of the T2-weighted anatomical images, was used to co-register the images to the b0 image of the diffusion data using the SPM12 toolbox (https://www.fil.ion.ucl.ac.uk/spm/software/spm12/). Within the white-matter mask, 200,000 most probable fibres were sampled for each participant. Then, fibres were transformed to the standard Montreal Neurological Institute (MNI) space applying a nonlinear deformation field derived from the T2-weighted images via a diffeomorphic registration algorithm^85^. The individual tractograms were then aggregated into a joint dataset in MNI standard space resulting in a normative tractogram representative of a healthy young adult population and made available in the leadDBS toolbox^84^.

The *NxN* matrices were computed from the normative tractogram using the Automated Anatomical Labelling (AAL) parcellation scheme^86^ with N=90 cortical and subcortical areas, by calculating the number of tracts, *C*(*n,p*), and mean tract length, *D*(*n,p*), between the voxels belonging to each pair of brain areas *n* and *p*. Further details on the structural matrices in the AAL and other parcellation schemes are reported in Supplementary Methods 1 and Supplementary Figure 14.

### MEG power spectra from healthy participants

The power spectra from human resting-state MEG signals were also downloaded from the Human connectome Project (HCP) database as a FieldTrip structure in a MATLAB file. The MEG power spectra are provided for 89 healthy participants at rest (mean age 28.7 years old, range 22–35, 41 female) distinct from the 32 participants from which the structural connectomes were derived, but with similar age range and gender ratio. Resting-state MEG signals were recorded on a Magnes 3600 MEG (4D NeuroImaging) with 248 magnetometers for 6 minutes and the “powavg” pipeline was used to obtain the power spectrum of the resting state MEG data in each MEG sensor. Briefly, the signals were segmented, Hanning-tapered, Fourier-transformed and the power spectrum was averaged over all segments. Notch filters were applied to remove the power line noise (cut-off frequencies 59-61 Hz and 119-121 Hz). Additional details are explained in the HCP reference manual (https://humanconnectome.org/storage/app/media/documentation/s1200/HCP_S1200_Release_Reference_Manual.pdf). The MEG power spectra were averaged across the 248 sensors to obtain a power spectrum representative of each subject.

### Brain Network Model

The Stuart-Landau (SL) equation (first term in Equation 1) is the canonical form to describe the behaviour of a system near an Andronov-Hopf bifurcation, i.e. exhibiting the birth of an oscillation from a fixed point^42,87^. In other words, it is used to describe systems that have a static fixed point (like a resting spring), but respond to perturbation with an oscillation, which may damped or self-sustained depending on the operating point of the system with respect to the bifurcation (see Supplementary Note 1 and Supplementary Figures 1-4). This model allows to describe complex-systems behaviour among several applications, bridging the gap between the simplicity of the Kuramoto model and the extensiveness of the phase-amplitude frameworks^88,89^. It describes how the oscillator behaves both when it is weakly attracted to a limit cycle (displaying only damped oscillations in response to perturbation) and, on the other hand, when it is purely restricted to a limit cycle (oscillations remain self-sustained).

Our analysis is based on a system of N=90 SL oscillators coupled in the connectome, considering both the connectivity strength, *C*_*np*_, and the conduction delays, *τ*_*np*_, between each pair of brain areas *n* and *p*. The conduction delays are defined in proportion to the fiber lengths between brain areas, assuming an homogenous conduction speed *v*, such that *τ*_*np*_ = *D*_*np*_ /*v*, where *D*_*np*_ is the length of the fibres detected between brain areas *n* and *p*. To simulate how the activity in node *n* is affected by the behaviour of all other nodes *p* (*p* ∈ *N* Λ *p* ≠ *n*), we describe the interaction between nodes in the form:

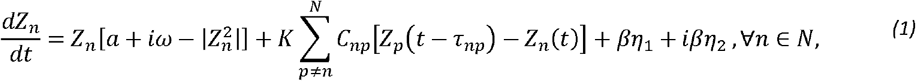

where the complex variable *z*_*n*_ (*t*) describes the state of the *n*^*th*^ oscillator at time t. The first term in equation (1) describes the intrinsic dynamics of each unit, the second term describes the input received from coupled units and the last terms represent uncorrelated white noise (see Supplementary Note 2 for detailed analysis of the model).

With this approach, we consider that the first term of Eq. 1 represents the natural excitability of neuronal assemblies, where ω = 2*π* * *f*_*f*_ is the angular frequency, with *f*_*f*_ as the fundamental frequency in Hertz. For our proof of concept, we set all nodes with identical natural frequency *ω*_0_ = 2*π* * 40*Hz*, representing the undifferentiated ability of a neural mass to engage in gamma-frequency oscillations.

The parameter *a* determines the position of the each unit with respect to the limit cycle. For *a* > 0, a stable limit cycle appears via a superciritical Hopf bifurcation, while when *a* < 0 there is only a stable fixed point at the origin *Z*_*n*_ = 0, so the bifurcation point is at *a* = 0. Importantly, if *a* is sufficiently close to the bifurcation, the system is still weakly attracted to the limit cycle and damped oscillations emerge in response to external input, with a decay time scaled by *a*. In this work, we pick a value of *a* = −5 for all nodes, such that a single input drives a damped oscillation decaying after ∼1s, approximating the slowest decay time-constants of inhibitory receptors (Supplementary Figure 4) (*τ*decay(GABAB) ≈ 500-1000ms). In Supplementary Note 2 and Supplementary Figure 7 we show that our results are qualitatively similar for a broad range of *a* values, both positive and negative, thus demonstrating the generalizability of synchronization at collective frequencies to coupled oscillatory systems with fluctuating amplitude, be they damped or self-sustained. We note that this mechanism only fails when the units have an overdamped response (exponential decay without oscillation), which, in this case, only occurred for *a* = −500. Thus it is of great interest in future research to investigate whether the local bifurcation parameters can be tuned based on sensitive observables to fit the MEG data of different individuals in different conditions.

The second term represents the total input received from other brain areas, scaled by parameter *K*, which sets the strength of all network interactions with respect to the intrinsic node dynamics. Because we wish to focus on the nonlinear phenomena introduced by time delays, we model the node-to-node interactions using a particular *linear diffusive coupling*, as the simplest approximation of the general coupling function, considering *delayed* interactions. Here, the signal of node *n* at time *t* is calculated with respect to the activity of all other nodes *p* at time *t*−*τ*_*np*_ (where *τ*_*np*_ is the time delay between *n* and *p*), scaled by the relative coupling strength given by *C*_*np*_.

The third term of Equation 1 represents the added uncorrelated noise to each unit (with real and imaginary components *η*_1_ and *η*_2_). In this analysis, the system is perturbed with uncorrelated white noise, where *η*_1_ and *η*_2_ are independently drawn from a Gaussian distribution with mean zero and standard deviation *β* = 0.001 (integrated as 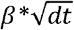).

In this framework, our whole-brain network model is purely bottom-up (i.e., not inferred from the MEG data we aim at explaining). For a qualitative comparison with the literature in delay-coupled oscillatory systems ^25,53,65^, we explore the network dynamics as a function of the coupling strength *K* and the mean delay ⟨*τ*⟩ = ⟨*D*⟩/*v*, where ⟨*D*⟩ is the mean length of all fibres detected between each pair of brain areas. For each set of parameters, the model is solved numerically for 50 seconds with an integration step *dt* = 10^−4^ seconds.

### Kuramoto Order Parameter

To evaluate the global synchrony of the simulated network activity over time, we use the Kuramoto order parameter:

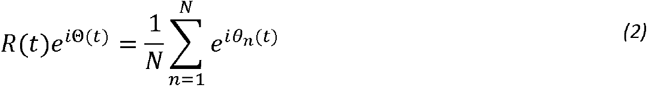

where *θ*_*n*_ (*t*) is the phase of each node, given by the argument of *Z*_*n*_. The temporal evolution of the Order Parameter *R*(*t*) provides an instantaneous measure of the degree of synchrony of the ensemble. Since we add noise in the simulations, we first band-pass filter the signals *Z*_*n*_ around the peak frequency of the ensemble. A steady order parameter indicates a stable solution (be it asynchronous, ⟨*R* (*t*)⟩ ∼0 or synchronous ⟨*R* (*t*)⟩ ∼1), whereas fluctuations in the order parameter are indicative of *Metastability*, driven by constant transitions between different weakly stable solutions^68^. For the analysis in parameter space, we take the mean ⟨*R* (*t*)⟩ as a measure of the global synchronization while the standard deviation *STD*(*R* (*t*)) indicates how much *R*(*t*) fluctuates in time^54^.

### Analytic Prediction of Collective Frequency of Synchronization

Previous theoretical studies have shown analytically that coupled oscillatory networks with homogeneous delays can find stable solutions at multiple collective frequencies Ω. Let us consider the Kuramoto transition in a population of phase oscillators defined by Equation 3:

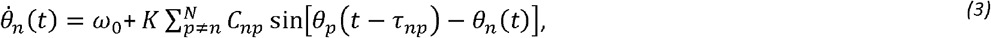

and the fully synchronized, uniformly rotating one-cluster state *θ*_*n*_ =… = *θ*_*N*_ = Ωt. Substituting this expression into Equation 3 we obtain ^25,26,65,81,90^:

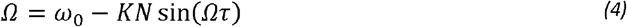

where ω_0_ corresponds to the nodes’ intrinsic frequency and *τ* is the homogeneous time delay between nodes. As *K* is increased and full synchrony is approached, the system finds an equilibrium point at the lowest stable solution for Equation 4, which is given by:

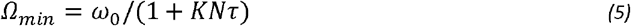

Note that, for collective oscillations to emerge, the global coupling *K* needs to be sufficiently strong such that the synchronized solutions are at least weakly stable. To approximate the analytic prediction from equation (5), the coupling matrix was normalized by its mean, such that <C>=1.

### Model Performance

We perform a parameter space exploration by tuning the two free parameters *K* and ⟨*τ*⟩. We choose to increase *K* exponentially as a power of 10 from 10^−1^ to 10^1.7^ in steps of 10^0.1^, to ensure a range that covers from weak to strong coupling. ⟨*τ*⟩ is explored in the range from 0 ms to 30 ms in steps of 1 ms.

We measure the fitting between the empirical sensor MEG PS for each of the 89 subjects and the simulated PS for each pair of parameters as the squared Euclidean distance, resulting in one fitting value for each subject. This can be regarded as a maximum likelihood procedure under the assumption of Gaussian observation noise.

### Metastable Oscillatory Modes

To detect MOMs and characterize them in space and time, we band-pass filter the simulated signals in each frequency band and obtain the corresponding amplitude envelopes using the Hilbert transform for each band. Subsequently, we consider that a node (or brain area) engages in a MOM if the amplitude increases 5 standard deviations above the amplitude in that frequency range. We define the baseline threshold considering the simulations with the optimal K but with zero delays. Since some areas are more coupled together than others, even with “zero delays” these areas may exhibit more power across frequencies that is purely due to noisy interactions. Therefore, we define a different threshold for each node and each band.

### Envelope Functional Connectivity Patterns

Following standard procedures to estimate frequency-specific functional connectivity in empirical source-projected MEG data^44^, we first bandpass filter the simulated signals in each frequency band of interest, compute the analytic signal using the Hilbert transform and then calculate the correlation matrices between the amplitude (i.e. the absolute value) of the analytic signal. This is done in one optimal point (K=10, ⟨*τ*⟩ =3ms), for weak coupling (K=0.1, ⟨*τ*⟩ =3ms), strong coupling (K=50, ⟨*τ*⟩ =3ms), no delays (K=10, ⟨*τ*⟩ =0ms) and long delays (K=10, ⟨*τ*⟩ =20ms). The same analysis performed using N=200 units is shown in Supplementary Methods 1 and Supplementary Figure 17.

## Data availability

Human neuroimaging data used in this study were provided by the Human Connectome Project (HCP)^83^ (https://www.humanconnectome.org), WU-Minn Consortium (Principal Investigators: David Van Essen and Kamil Ugurbil; 1U54MH091657) funded by the 16 NIH Institutes and Centers that support the NIH Blueprint for Neuroscience Research; and by the McDonnell Center for Systems Neuroscience at Washington University.

The normative connectomes were computed from Human Connectome Project data and included as part of the leadDBS toolbox^84^ (https://www.lead-dbs.org/).

The matrices computed from the normative connectomes used for simulations, together with the MEG power spectra from 89 individuals, are publicly available in .mat format at: https://github.com/fcast7/Hopf_Delay_Toolbox.

Simulated data is available from the corresponding author on reasonable request.

Supplementary Notes, Supplementary Methods and Supplementary Video 1 are available with this manuscript.

## Code availability

All simulations were performed in MATLAB2021b. The codes used in this study are publicly available at: https://github.com/fcast7/Hopf_Delay_Toolbox.

## Author contributions

JC, GD and MLK conceived and designed the analysis. JC, FC, JV, GD contributed with data and analysis tools. JC and FC performed the simulations and analysis. VL, KF, RL, CB, MLK and GD supervised the analysis. JC wrote the original draft. FC wrote the Supplementary Information. All authors reviewed and edited the final manuscript and Supplementary Information.

## Acknowledgments

JC is funded by the Portuguese Foundation for Science and Technology grants UIDB/50026/2020, UIDP/50026/2020 and CEECIND/03325/2017, Portugal. FC is funded by the EU-project euSNN European School of Network Neuroscience (MSCA-ITN-ETN H2020-860563). The Wellcome Centre for Human Neuroimaging is supported by core funding from Wellcome [203147/Z/16/Z]. JV is supported by the EU H2020 FET Proactive project Neurotwin grant agreement no. 101017716. RL acknowledges support from EPRSC Grants No. EP/V013068/1 and EP/V03474X/1. CB acknowledges support from the Engineering and Physical Sciences Research Council (EPSRC) through the grant EP/T013613/1. MLK is supported by the Center for Music in the Brain, funded by the Danish National Research Foundation (DNRF117), and the Centre for Eudaimonia and Human Flourishing, funded by the Pettit Foundation and Carlsberg Foundation. GD is supported by the Spanish national research project (PID2019-105772GB-I00 MCIU AEI), funded by the Spanish Ministry of Science, Innovation and Universities (MCIU), State Research Agency (AEI); HBP SGA3 Human Brain Project Specific grant agreement 3 (945539), funded by the EU H2020 FET Flagship program; SGR Research Support Group (reference 2017 SGR 1545), funded by the Catalan Agency for Management of University and Research Grants (AGAUR); Neurotwin Digital twins for model-driven non-invasive electrical brain stimulation (grant agreement 101017716), funded by the EU H2020 FET Proactive program; euSNN (grant agreement 860563), funded by the EU H2020 MSCA-ITN Innovative Training Networks; The Emerging Human Brain Cluster (CECH) (001-P-001682) within the framework of the European Research Development Fund Operational Program of Catalonia 2014–2020; Brain-Connects: Brain Connectivity during Stroke Recovery and Rehabilitation (201725.33), funded by the Fundacio La Marato TV3; and Corticity, FLAG-ERA JTC 2017 (reference PCI2018-092891), funded by the MCIU, AEI.

