## Supplementary Methods for "Metastable Oscillatory Modes emerge from synchronization in the Brain Spacetime Connectome"

### Supplementary Information

#### Supplementary Note 1: Uncoupled oscillator and intrinsic node dynamics

Approaching the neuronal ensemble in each brain area as an autonomous system displaying oscillations in response to an external input, the node dynamics can be described from a dynamical system's perspective as the birth of an oscillation from an equilibrium point. In mathematical terms, this phenomenon can be represented by the canonical form of a supercritical Andronov-Hopf bifurcation, also called a Stuart-Landau (SL) oscillator, which is the model for investigating the appearance of an oscillatory mean field from a noisy interacting units [1,2].

Nodes in a brain network represent brain areas composed of millions of densely interconnected neurons organized in cortical columns. Even though some brain areas are responsible of highly specialized functions, they all have a property in common: they are composed of excitatory and inhibitory neurons whose ensemble activity can generate coherent voltage fluctuations in the local field potential (LFP) between $10^{-4}$ to $10^{-3}$ Volts, even when spiking neurons display irregular action potentials at different firing rates [3-7]. This intrinsic rhythmogenesis of neural masses has been induced *in vitro* in isolated cortical and subcortical slices in the gamma frequency range (35-80Hz, where 40Hz is typical) [3,4,8,9] and has been linked to feedback inhibition within the neuronal ensemble [10-13]. For the purpose of dimensionality reduction in this work, we take the intrinsic oscillatory frequency of LFPs in the gamma-frequency range (where 40Hz is typical) as a key ingredient of our network model bridging the activity at the neuronal level with the mesoscopic dynamics of the neuronal ensemble at each node.

For now, let us consider the dynamics of a single unit (uncoupled node *n*), given by the following ordinary differential equation in complex Cartesian coordinates:

$$\frac{dZ}{dt} = Z[a + i\omega - |Z^2|] + \beta\eta_1 + i\beta\eta_2 \tag{S1}$$

where $Z = x + iy$, with *i* representing an imaginary unit. In this formula, $a$ is the bifurcation parameter and $\omega = 2\pi * f_f$ is the angular frequency, where $f_f$ is the intrinsic oscillatory frequency. When $Z$ is shifted by an external perturbation, the system responds with an oscillation with angular frequency $\omega$, which may be damped or self-sustained depending on the sign of the constant $a$. Generally speaking, each uncoupled oscillator has a supercritical bifurcation at $a = 0$, corresponding to the *isochrono*us state when phases and amplitudes are untangled. On the other hand, if $a \neq 0$, the oscillations appear to be *non-isochronous*, that is, the frequency depends on the amplitude.

Different regimes can be obtained by simply varying the local parameter $a$, which determines the operating point of the system with respect to the bifurcation:

- <u>a>0: *supercritical regime:*</u> when the parameter $a$ is positive, the system exhibits a periodic oscillation, with fundamental frequency $f_f = \omega/2\pi$, which corresponds to a closed curve in the phase plane, named *limit cycle.* This behaviour is typical of self-sustained oscillators: after perturbing their oscillation, the natural rhythm is restored. This is also called a *simple attractor*, in contrast to the concept of *strange attractor* where oscillators exhibit a *chaotic* motion (Supplementary Figure 1, Right).
- <u>a<0: *damped subcritical regime:*</u> when the parameter $a$ is negative, the system decays to a fixed point equilibrium upon perturbation. If $a$ is sufficiently close to the bifurcation (and this range depends on the natural frequency $\omega$) the system exhibits damped oscillations with decaying amplitude (Supplementary Figure 1, Middle). The more $a$ is negative, the faster the decay, such that beyond a certain value of negative $a$, the system decays without any oscillation, and is said to be *overdamped (*Supplementary Figure 1, Left).

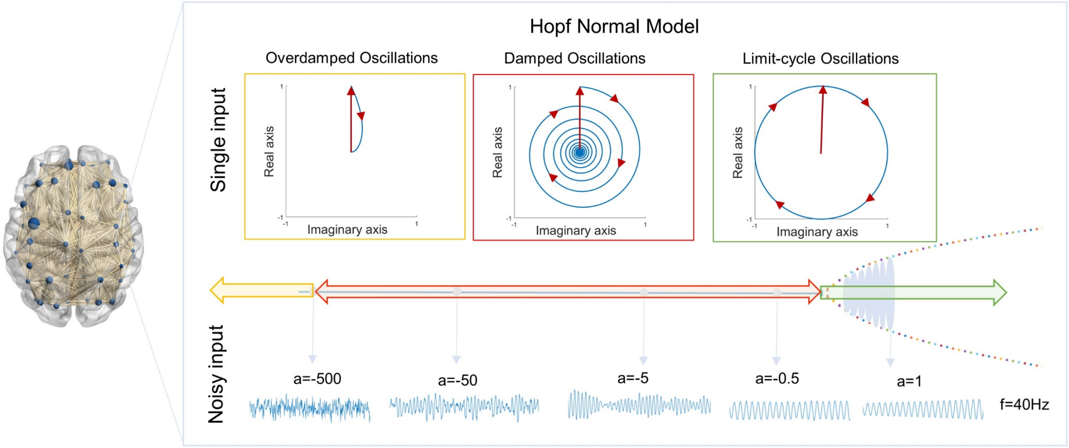


**Supplementary Figure 1. Andronov-Hopf Normal Form. Uncoupled node response to a noisy Gaussian input with different values of bifurcation parameter '*a*' and an intrinsic frequency of 40Hz.** (Top) Each dynamical oscillatory unit, responding to a single perturbation with an oscillation at 40Hz whose amplitude: **left -** decays exponentially toward the equilibrium fixed point at the origin without oscillating; **middle -** decays slower toward to an equilibrium point exhibiting one or more sustained cycles; **right-** remains constant (limit-cycle oscillator). (Bottom) For modelling local neural populations we used the normal form of a Hopf bifurcation perturbed with noise where, depending on the value of the bifurcation parameter, the local model generates a chaotic signal (left), oscillations with fluctuating amplitude (middle) or a pure oscillatory signal (right).

An oscillator cannot be considered as being completely isolated from the environment; therefore, it is subject to thermal fluctuations and its behaviour consists of decaying, free oscillations depending on the initial conditions (the *homogeneous* part) plus forced oscillations depending on noise only (the *inhomogeneous* part). But how does noise affect an autonomous oscillator? Even a weak noise is known to affect the phase, shifting the signal back and forth. To illustrate the effect of a random perturbation to an uncoupled node, we add complex Gaussian white noise, where $\eta_1$ and $\eta_2$ are independently drawn from a Gaussian distribution with mean zero and standard deviation $\beta = 0.001$. We use the Euler integration method, and the noise term is multiplied by $\sqrt{dt}$ so that $noise = \beta * \sqrt{dt}$ (Supplementary Figure 1, Bottom).

More specifically, when the system is operating in the subcritical regime but sufficiently near the bifurcation to exhibit damped oscillations, noisy perturbations induce the emergence of damped oscillations at the fundamental frequency $f_f$ leading to a peak in the power spectrum at the corresponding frequency.

To justify the choice of the bifurcation parameter $a$, we define how close to the bifurcation point the oscillator must operate. It is in the damped subcritical regime that the simulated signals better resemble the empirical neural recordings [14]. However, the range of $a$ values for which damped oscillations occur depends on the intrinsic frequency: signals with high intrinsic frequency (i.e. LFPs, $\omega_{hf} = 2\pi * 40Hz$) exhibit damped oscillations for a larger range of $a$ values compared to those with lower intrinsic frequency (i.e. fMRI signals, $\omega_{lf} = 2\pi * 0.05Hz$), where the range is shorter and $a$ is commonly selected close to zero (i.e., simulations of ultra-slow fMRI activity typically use values of $a$ between -0.5 and 0.5 [14,15]).

Therefore, for the same '$a$' (i.e., $a$ =-5 in Supplementary Figure 2), a system with a high fundamental frequency (here, $f_f$=40Hz) may exhibit damped oscillations (leading to a distinct peak in the power spectrum at 40Hz), whereas for a system with a lower fundamental frequency, the oscillations at the peak frequency less distinguishable from noise.

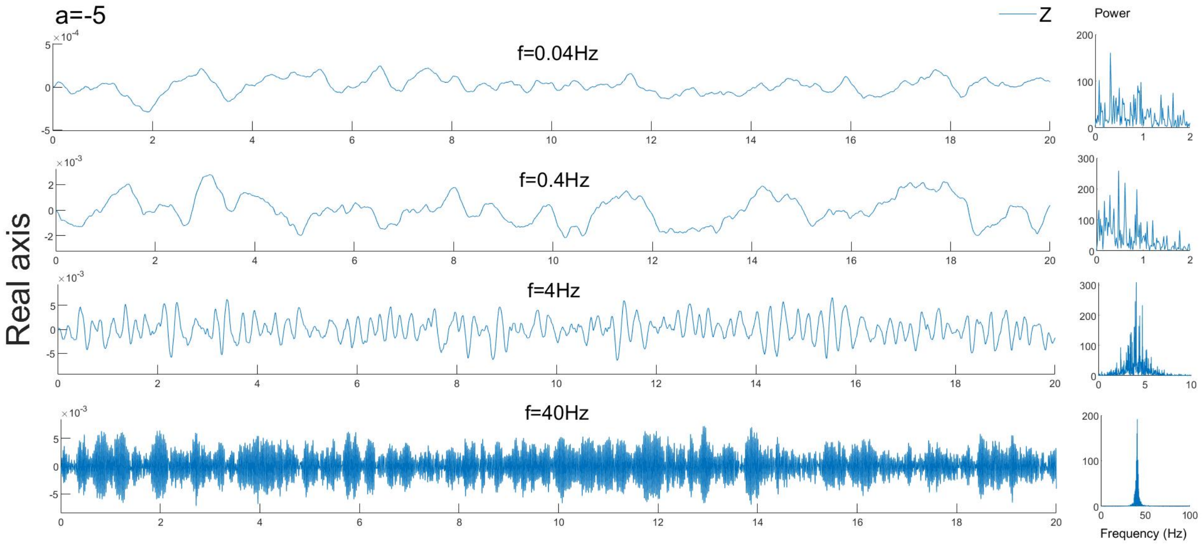


**Supplementary Figure 2. Stuart-Landau oscillators with different intrinsic frequencies 'f' and fixed bifurcation parameter '$a$'.** **Left -** Simulated signals (taking the real part of Z) in the damped regime with $a$ =-5 and different intrinsic frequencies, perturbed with complex Gaussian white noise. **Right -** Power Spectrum of the signals simulated over 50 seconds.

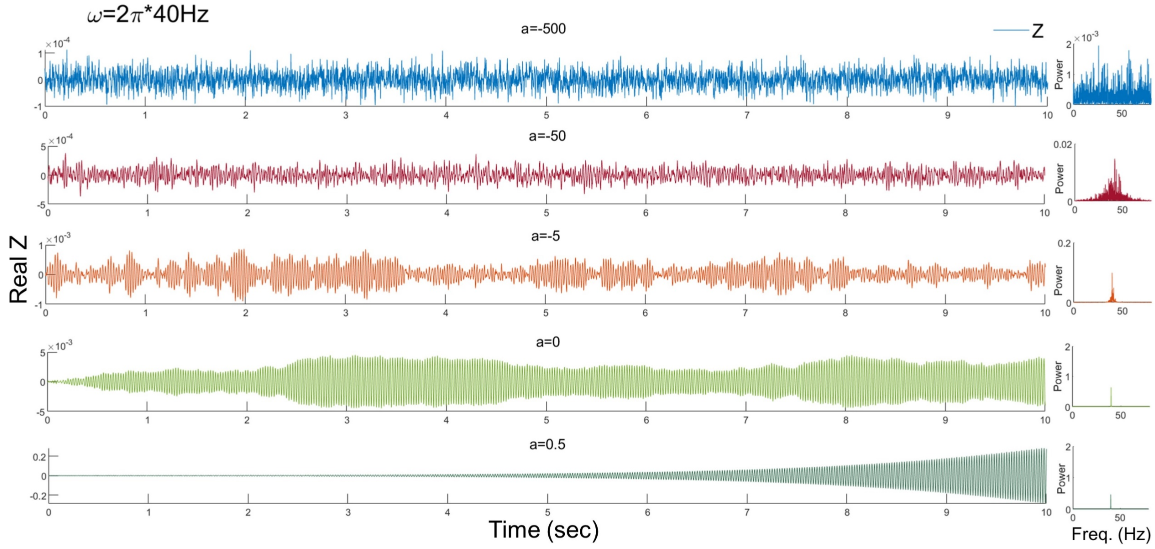


**Supplementary Figure 3. Stuart-Landau oscillators in the presence of Gaussian white noise with different values of bifurcation parameter '$\boldsymbol{a}$' and fundamental frequency of 40Hz. Left -** Numerical simulations of a Hopf unit with intrinsic frequency ω=40Hz, in response to persistent input in the form of Gaussian white noise. In the presence of a constant noisy input, the system displays oscillations with time-varying amplitude depending on the value of the bifurcation parameter $\boldsymbol{a}$. We find that the resonant peak at the fundamental frequency only disappears for $\boldsymbol{a}$ =-500. **Right -** Power Spectrum of the signals simulated over 50 seconds.

We want $a$ to be in a regime where noise induces gamma oscillations with fluctuating amplitude[7]. Therefore, as a first point of our analysis, we need to tune $a$ to be somewhere below the limit-cycle regime but above the overdamped regime where the oscillatory response is lost (pure noise). For this project, we pick a value of $a = -5$ for all nodes, such that a stochastic input drives a damped oscillation decaying after ~1s, approximating the slowest decay time-constants of inhibitory receptors (Supplementary Figure 4) ($\tau_{decay}$($GABA_B$) ≈ 500-1000ms). However, the choice of $a$ goes beyond the dynamics of a single unit: uncoupled and coupled scenario may exhibit two different behaviours, considering how complex the interactions are between oscillators. Accordingly, a whole-brain parameter space exploration and optimization should be considered for more detailed analysis, for example by measuring the Euclidean distance between Model Functional Connectivity and Empirical Functional Connectivity for different combination of the global coupling strength, the bifurcation parameter, and time delay.

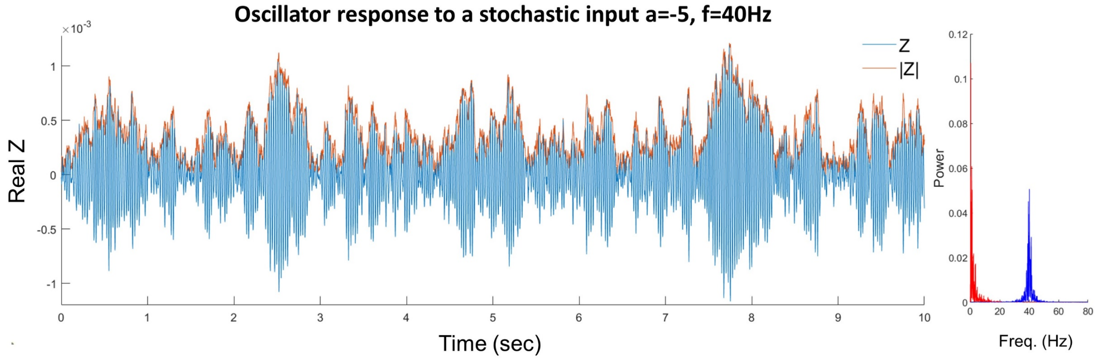


**Supplementary Figure 4. Stuart-Landau oscillator response to a stochastic Gaussian input with 'a'=-5 and fundamental frequency of 40Hz. Right -** The intrinsic resonance of local field potentials in the gamma frequency is represented by the real part of Z in the damped regime with $\boldsymbol{a}$=-5 and natural frequency f=40Hz. Z is perturbed with complex Gaussian white noise.

**Left -** Power Spectrum of the Phase and Amplitude dynamics (blue and red). After perturbation, the phase of Z oscillates at the fundamental frequency of 40Hz, whereas the amplitude of Z reveals slow fluctuations with 1/f spectrum.

In summary, we selected the bifurcation parameter to obtain a fair approximation of the local field potential of neural masses for the bottom-up model. However, as we will show in section II, the model results remain qualitatively similar for different node regimes, as long as the system is not overdamped and oscillations emerge (i.e., for a>-500).

In previous works from our group, the phenomenological case of coupled self-sustained oscillators was addressed using the Kuramoto model with time delays [16,17]. However, since the inspiring paper by Rosenblum, Pikovsky and Kurths, it is well recognized that amplitude affects synchronization [18]. While phase-only models such as the Kuramoto model have been extensively applied to study synchronization in brain oscillations, phase-amplitude models are shown to capture richer behaviours than phase-only ones [19,20].

#### Supplementary Note 2: Coupled Stuart-Landau Units

Let us consider a population of identical SL oscillators in the presence of noise:

$$\frac{dZ_n}{dt} = Z_n[a + i\omega - |Z_n^2|] + \sigma \sum_{p \neq n}^{N} C_{np}\left[Z_p\left(t - \tau_{np}\right) - Z_n(t)\right] + \beta\eta_1 + i\beta\eta_2, \qquad \text{(S2)}$$

where the complex variable $\frac{dZ_n}{dt}$ describes the state of the n[th] oscillator; $\sigma = Ke^{i\varphi}$ is the complex coupling strength and $C$ is the real-valued symmetric connectivity matrix. We describe the behaviour for the coupling phase $\varphi = 0$, where the collective frequency is distributed around the intrinsic frequency $\omega = 40Hz$.

The final equation for the modelling exercise in complex coordinates is therefore:

$$\frac{dZ_n}{dt} = Z_n[a + i\omega - |Z_n^2|] + K \sum_{p \neq n}^{N} C_{np}\left[Z_p\left(t - \tau_{np}\right) - Z_n(t)\right] + \beta\eta_1 + i\beta\eta_2 \qquad \text{(S3)}$$

And, given that $z = re^{i\theta} = x + iy$, equation S3 becomes in polar coordinates

$$\dot{r}_n(t) = r_n(t)[a - |r_n^2|\,] + K \textstyle\sum_{p \neq n}^{N} C_{np} r_p cos\left[\theta_p\left(t - \tau_{np}\right) - \theta_n(t)\right] \qquad \text{(S3.1)}$$

$$\dot{\theta}_n(t) = \omega + K \textstyle\sum_{p \neq n}^{N} C_{np} \frac{r_p}{r_n} sin\left[\theta_p\left(t - \tau_{np}\right) - \theta_n(t)\right], n = 1,2 \ldots N \qquad \text{(S3.2)}$$

where $r_n(t)$ and $\theta_n(t)$ are the amplitude and phase of the signal, respectively, produced by the oscillator at time $t$. For a detailed exploration and dynamical analysis of SL model see [21,22].

*The coupling strength and time delays*

With delays, the brain network model is described mathematically by a set of coupled delay differential equations. Time delays have been shown to profoundly affect the synchronization dynamics of chaotic coupled oscillators, with the ability of anticipate and control their time evolution [23-25]. This is the case especially if they are in the same order of magnitude of the natural period of intrinsic oscillations[1,26,27]. Noticeably, while for null delays one synchronization state is stable, for nonzero delays and a given value of K, *Multistability*[1] does take place (i.e., there are multiple state solutions for Equation S2) with multiple basins of attraction and synchronization frequencies.

*Stuart-Landau to Kuramoto*

The advantage of reducing the complexity of the brain network to a simple phenomenological model is that it allows analysing our results in the light of *universal principles* governing the dynamics of delay-coupled nonlinear systems. Still, most of the literature in this domain has been focused in coupled phase oscillators using the Kuramoto model. From exploration of self-emerging complex structure with time delays [28-30] to investigation within multidimensional space [25], this model has proven to explain a variety of interesting behaviours, mirroring the most complex system in nature, but the way it extends to phase-amplitude models remains mostly unexplored. In their work, Gambuzza and colleagues have shown that when the bifurcation parameter is larger compared to coupling K, the amplitude dynamics of a SL oscillator vanishes, i.e., $\dot{r}_n(t) = 0$ and $r_n(t) = \sqrt{a}$ (constrained to its value on the limit cycle) and equation (2) can be reduced to:

$$\dot{\theta}_n(t) = \omega + K \sum_{p\neq n}^{N} C_{np} sin\left[\theta_p\left(t - \tau_{np}\right) - \theta_n(t)\right] \qquad \text{(S4)}$$

which is the Kuramoto model for in a population of coupled phase oscillators with time delays[31]. However, in this work, for the sake of biological plausibility, we choose a negative $a$, so this equivalence cannot be applied. Nevertheless, we hypothesize that the phenomenon of synchronization at reduced collective frequencies described for delay-coupled phase oscillators can extend to delay-coupled subcritical phase-amplitude models.

*Spectral properties of the network: emergence of collective frequencies*

In their remarkable application in 1999, Yeung and Strogatz [23] showed that coupled oscillators with homogeneous time delays can find solutions where both the incoherent and the synchronized states are stable (bi-stability). In 2009, Lee, et al. [24] extended the analysis to heterogeneous delays and reported a range of parameters supporting multi-stability, where a number of coherent attractor states can coexist due to weakly-stable cluster synchronization at the border between incoherence and full synchrony[24].

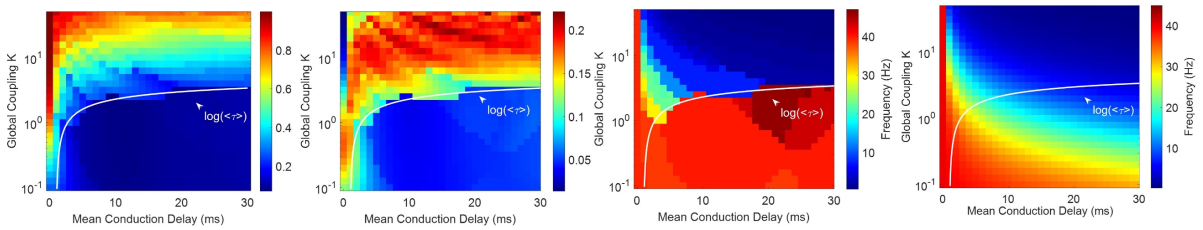


Supplementary Figure 5. The critical of K between incoherence and synchrony is found to depend logarithmically on the mean conduction delay.

Relatedly, our analysis shows how the spectral signature of the network is modulated by the global coupling strength between nodes. For low inter-node coupling ($K = 0.1$) , all nodes display only locally-generated rhythms at $\omega_0$=40Hz (Supplementary Figure 6). As the coupling strength increases

[1] Many different synchronous states as well as several stable phase configurations may be possible when two oscillators interact

within the range $0.1 < K < 2$, the power spectrum becomes broader, but still centred at 40Hz, indicating that the delayed node-to-node interactions are only sufficiently strong to induce small deviations to the unit's intrinsic 40 Hz limit-cycle (as occurring purely in the presence of white noise, as shown in Supplementary Figure 2). For $K > 2$ the network dynamics crosses a critical border and 40Hz oscillations become unstable giving place to a broad range of frequencies covering the whole spectrum but peaking at a slow frequency approximating the slowest collective frequency predicted analytically for delay-coupled phase oscillators. As the coupling increases further ($K \geq 2$) the system becomes more synchronized, the power at low frequencies is increased, while the power at high frequencies is suppressed. Due to the natural heterogeneity of the system and to the added noise, the fully synchronized solution of the system is unstable even at $K \approx 50.1$.

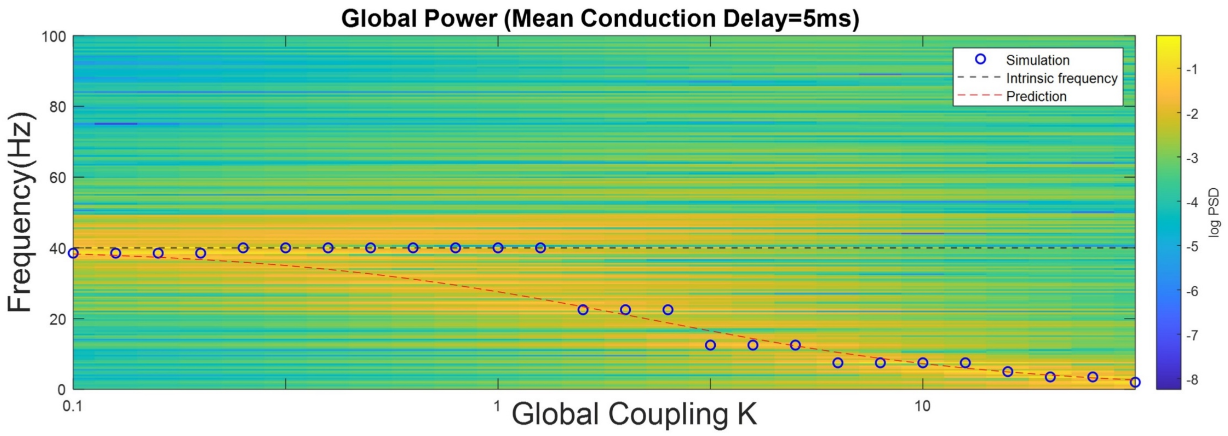


Supplementary Figure 6. Collective synchronization occurs at K-dependent reduced frequencies. Fixing the transmission speed between nodes (with a mean delay of $\langle\boldsymbol{\tau}\rangle$=5ms) we explore the influence of the global coupling strength K in the frequency spectrum between 0 and 100Hz of the simulated signals in all N=90 nodes, $\boldsymbol{Z_n}$. For weak coupling (K=0.1) all nodes oscillate close to their natural frequency $\boldsymbol{\omega_0}$=40Hz whereas for high coupling ($\boldsymbol{K \approx 50}$) all nodes oscillate around the collective frequency $\boldsymbol{\Omega_{min}}$=0.5Hz predicted by equation 8 (red dotted line).

Another relevant fact to take into account is that the presence of time delays in oscillatory systems facilitates what is termed in Physics *amplitude death* [32]. In other words, collective synchronization induces not only a shift in the peak frequency of the oscillators, but the stability of the collective ensemble is accompanied by a substantial suppression of the oscillations' amplitude [33]. Delay can cause transitions from amplitude death to periodic oscillations via Hopf bifurcations, growing or reducing the delayed limit cycle as we shift the delay towards or away from the critical bifurcation point [34].

Additionally, Han and colleagues [35] have also investigated the multistability phenomenon in locally complex conjugate coupled Stuart–Landau oscillators, remarking the importance of *amplitude death*. Corroborating their previous findings [36], they found that the stability of the amplitude death is independent of the number of oscillators. On the other hand, when the amplitude death state is unstable, a large number of states such as homogeneous oscillation death, heterogeneous oscillation death, homogeneous oscillations, and wave propagations emerge, and they may coexist.

This recently described phenomenon has high implications not only for the interpretations of our simulation results (since the bifurcation parameter in the Hopf model scales the amplitude, influencing the system's dynamics) but also for understanding the discrepancies in power spectra between intracranial LFPs and M/EEG signals.

*Effect of the bifurcation parameter in the network dynamics*

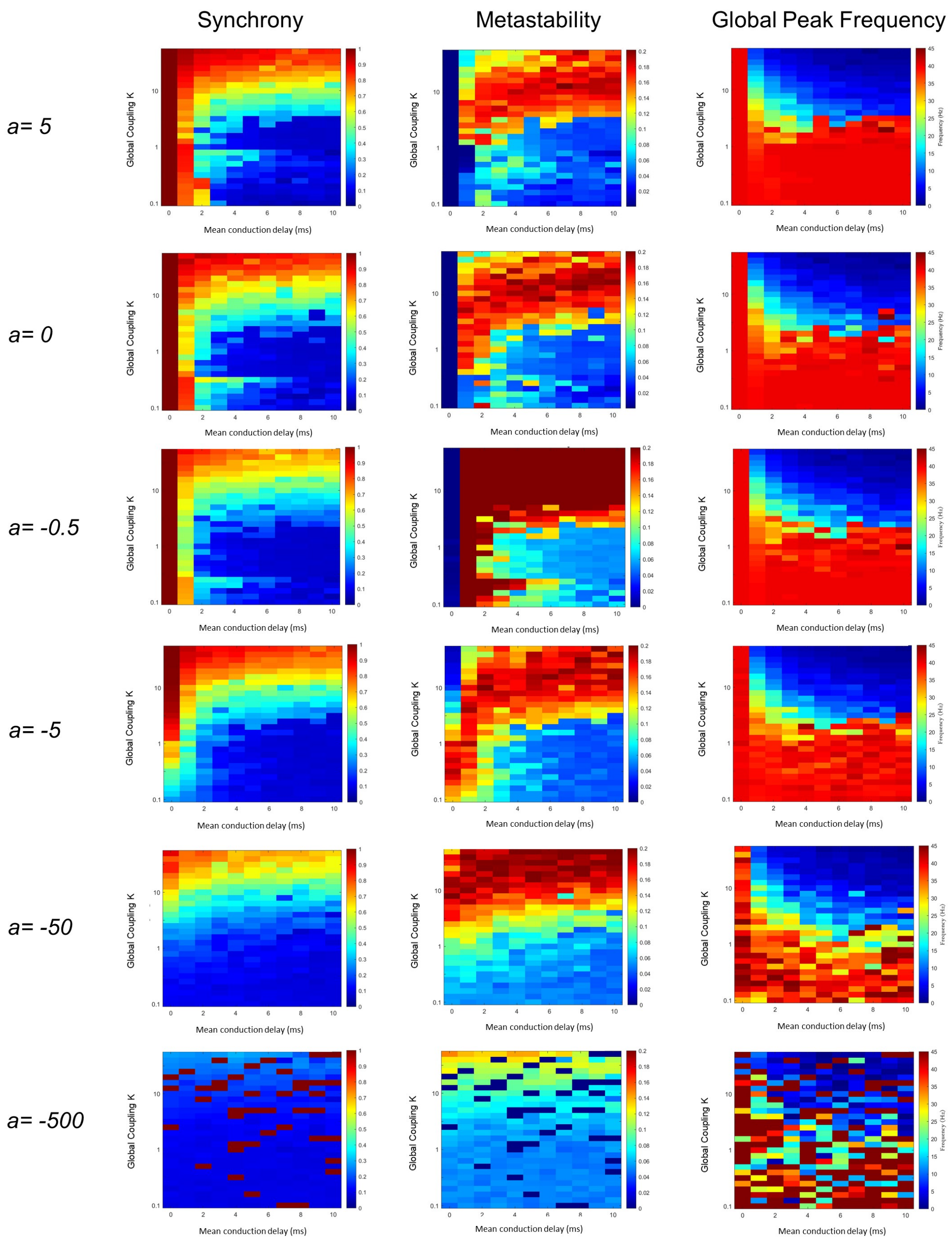


**Supplementary Figure 7. Synchronization at reduced frequencies is observed not only for oscillators in the limit cycle regime (supercritical, a>0), but extends to underdamped oscillators (subcritical).** The mean and standard deviation of the order parameter (referred to as Synchrony and Metastability accordingly) and the peak frequency of the global signal (i.e., summing all N=90 simulated signals), are reported for the range of couplings and delays explored for different values of the bifurcation parameter.

#### Effect of frequency dispersion on the Kuramoto Order Parameter

a-

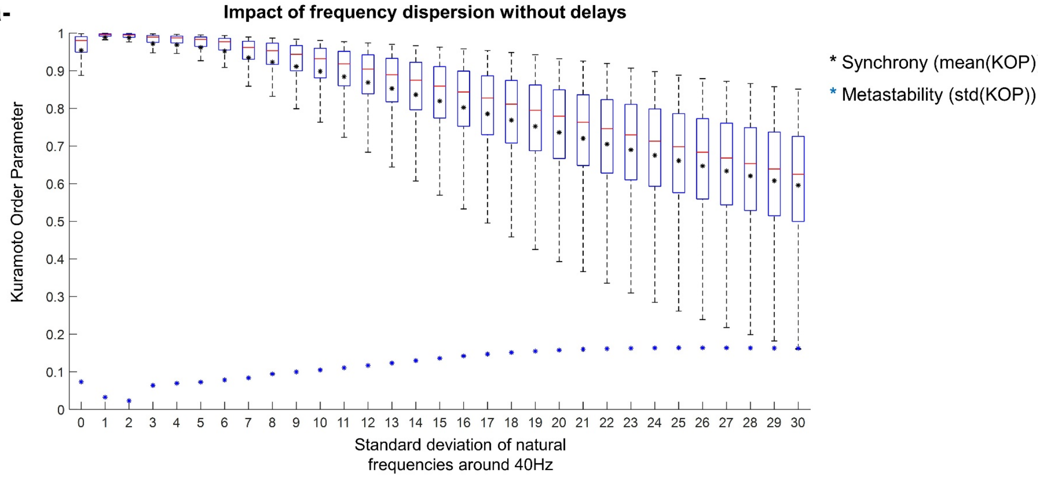


b-

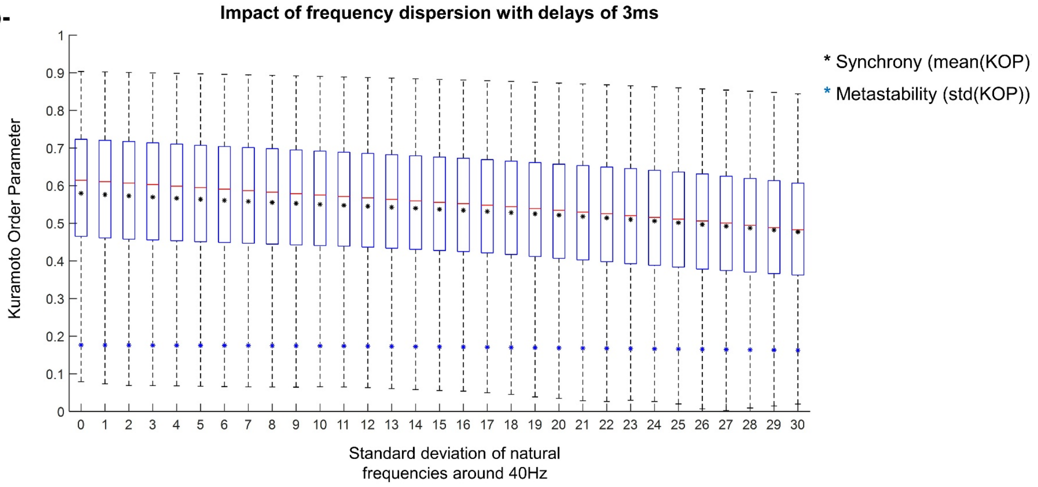


**Supplementary Figure 8. The Kuramoto order parameter as a function of the frequency dispersion for N=90 nodes in one optimal point.** (Top) Impact of frequency dispersion without delays. (Bottom) Impact of frequency dispersion with delays of 3ms. The Kuramoto Order Parameter (KOP) is evaluated across N=90 nodes for values of the standard deviation of the natural frequencies around 40Hz ($std(\omega_{40Hz})$ ranging from 0Hz to 30Hz in steps of 1Hz). Synchrony (mean(KOP)) and metastability (std(KOP)) are reported as black and blue asterisks, respectively. Generally, the KOP value decreases as the frequency dispersion increases. a- when delays are not considered (MD=0 with a fixed K=10), the frequency dispersion has a stronger influence on the KOP: the synchrony value goes from 1 to approximately 0.6, which shows that the system goes from a fully synchronised state to a weakly synchronized one. On the other hand, the metastability increases, showing that there are no fluctuations in the KOP for low values of $std(\omega_{40Hz})$ but they start to emerge as the $std(\omega_{40Hz})$ increases. b- When delays are considered (MD=3ms, with a fixed K=10), the impact of the frequency dispersion is less visible when compared to the delayed profile. Despite the decrease in synchrony values, there is no state-switch: the system is weakly synchronized for all $std(\omega_{40Hz})$ values. The metastability values remain approximately constant around 0.18, which is indicative of high metastability in the system no matter how the natural frequencies disperse.

*Power spectra of empirical MEG signals*

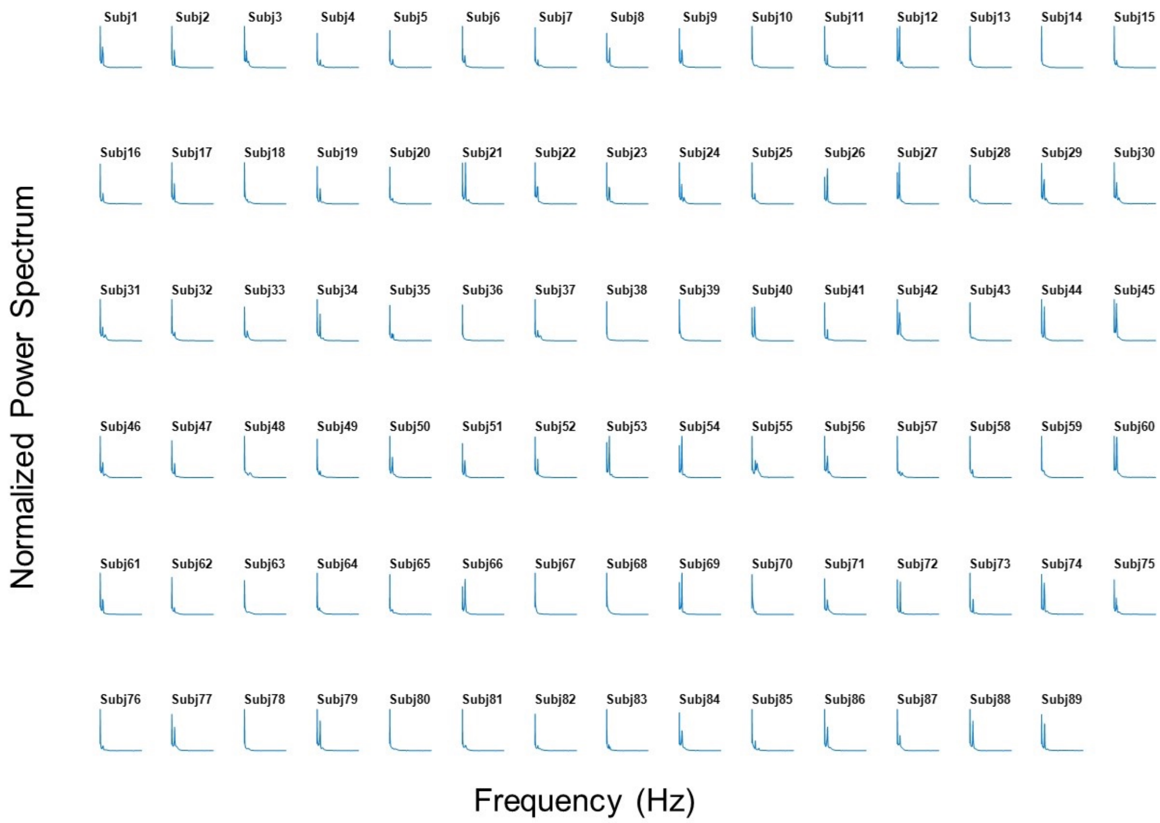


**Supplementary Figure 9 - The normalised power spectrum for each of the 89 healthy individuals from the Human Connectome Project open-source database.** The power spectra provided for each MEG sensor between 0 and 80 Hz was averaged across sensors to obtain one 'global' power spectrum representative of each subject.

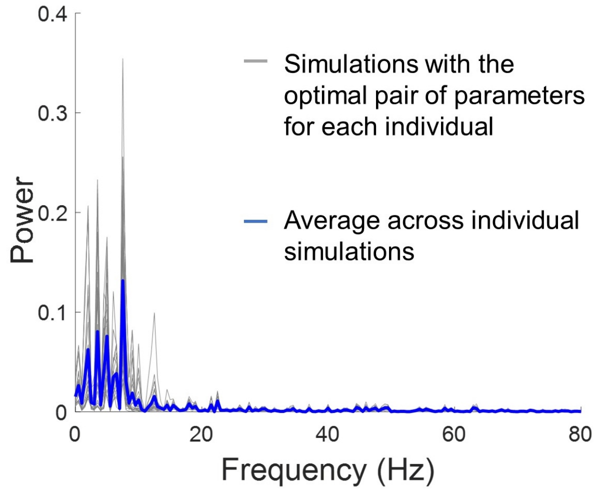


**Supplementary Figure 10 - Power Spectrum of the simulated signals for the pairs of free parameters that optimally approximated the MEG power spectra of healthy awake individuals.**

#### Supplementary Note 3: Metastable Oscillatory Modes

When the network model operates in a regime of high metastability, fluctuations in the magnitude of the order parameter are driven by metastable cluster synchronization, a phenomenon observed in systems of coupled oscillators with community structure at the transition between incoherence and synchrony[37,38]. In other words, when the coupling is strong, but not sufficiently strong to induce stability of the fully synchronized mode, some subsets of brain areas that are more strongly connected together (i.e., clusters/communities) can engage in partially-synchronized modes that remain stable for a given period in time but are naturally disrupted due to the heterogeneity in the system. Importantly, due to the presence of time delays, these coalitions do not synchronize at the natural frequency $\omega_0$ of the individual units, but instead oscillate at slower collective frequencies $\Omega$ determined by 1) the number of areas involved, 2) the time delays between them and 3) the strength of the coupling, as predicted by the Niebur equation[25]. Because of this mechanism, all areas engaging in a given coalition display the simultaneous emergence of an oscillation at the same collective frequency, whereas all other brain areas remain with low power at that frequency. This simple mechanism generates oscillations at the system level, which are organized in space (depending on the group of areas engaged in the coalition) and in time (depending on the relative stability of the coalition).

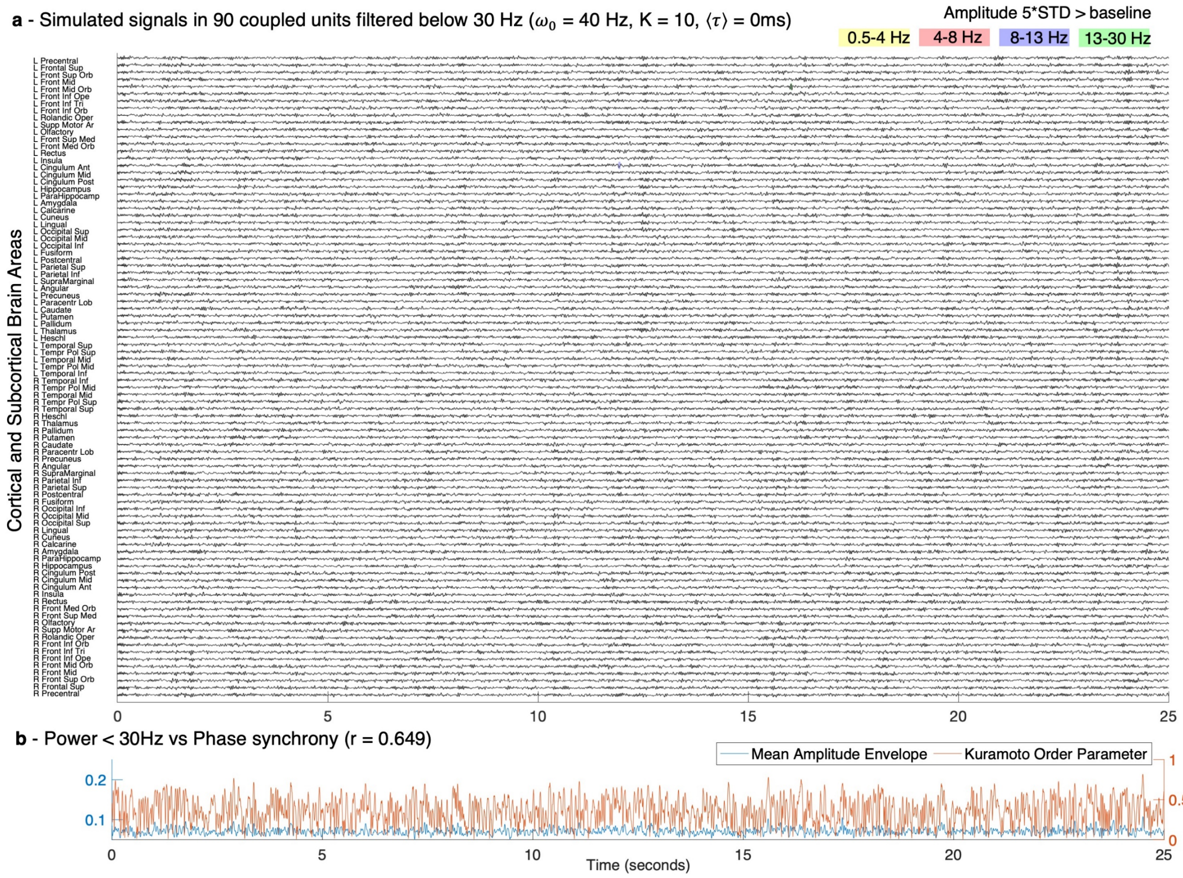


**Supplementary Figure 11. Baseline simulations. Time series for all 90 brain areas over 25 seconds without delays and intermediate coupling.** – The simulated signals over 25 seconds in all 90 units, each representing a brain area from the brain parcellation template, filtered below 30 Hz in order to concentrate on the sub-gamma oscillatory activity detected with most power with magnetoencephalography (MEG).

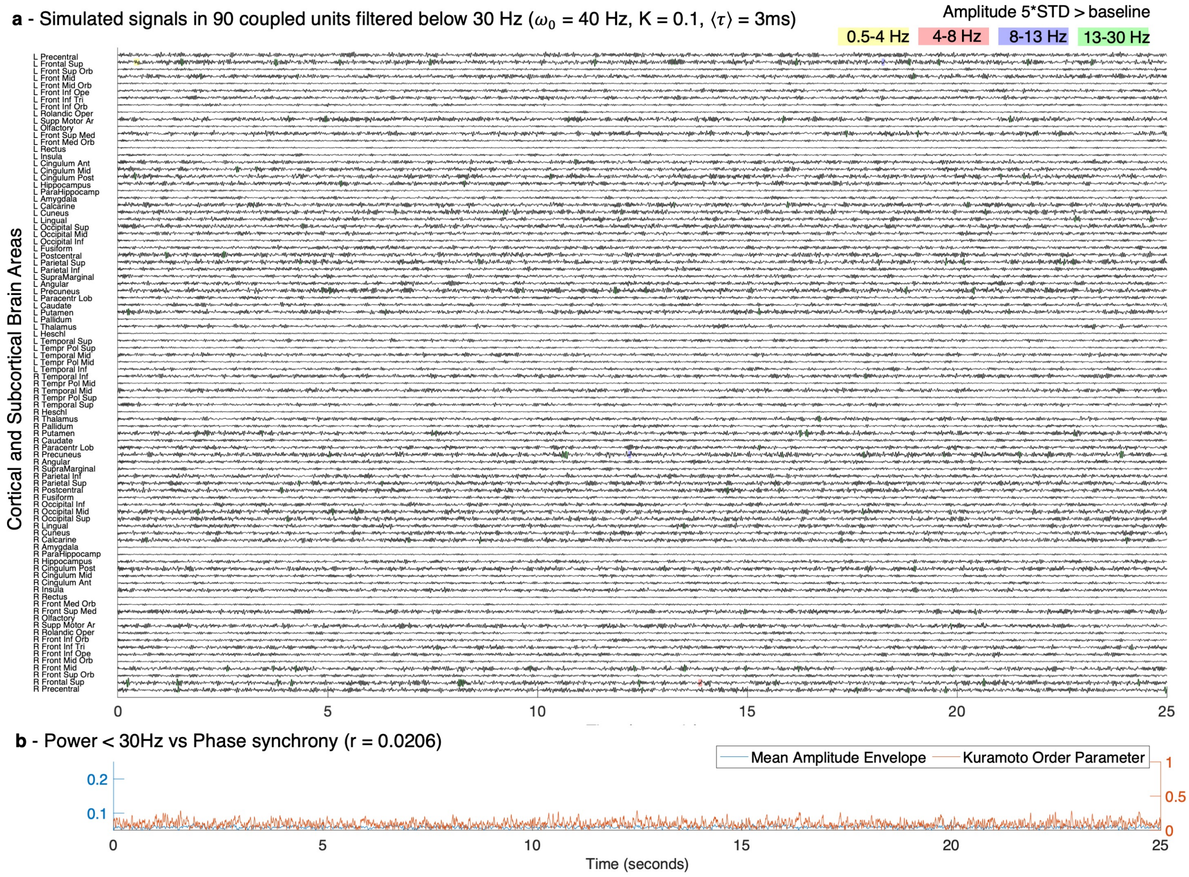


**Supplementary Figure 12. For weak coupling, no Metastable Oscillatory Modes (MOMs) are detected above threshold.** The simulated signals over 25 seconds in all 90 units, each representing a brain area from the brain parcellation template, filtered below 30 Hz in order to concentrate on the sub-gamma oscillatory activity detected with MEG. Shades highlight the time points of increased power in the delta (yellow), theta (red), alpha (blue) and beta (green) frequency bands, which in this case do not emerge due to the weaknesses of the coupling strength. For each frequency band, the threshold was defined as 5 standard deviations (STD) from the amplitude in the same frequency bands of the simulated signals when no delays were considered (baseline simulations Supplementary Figure 11).

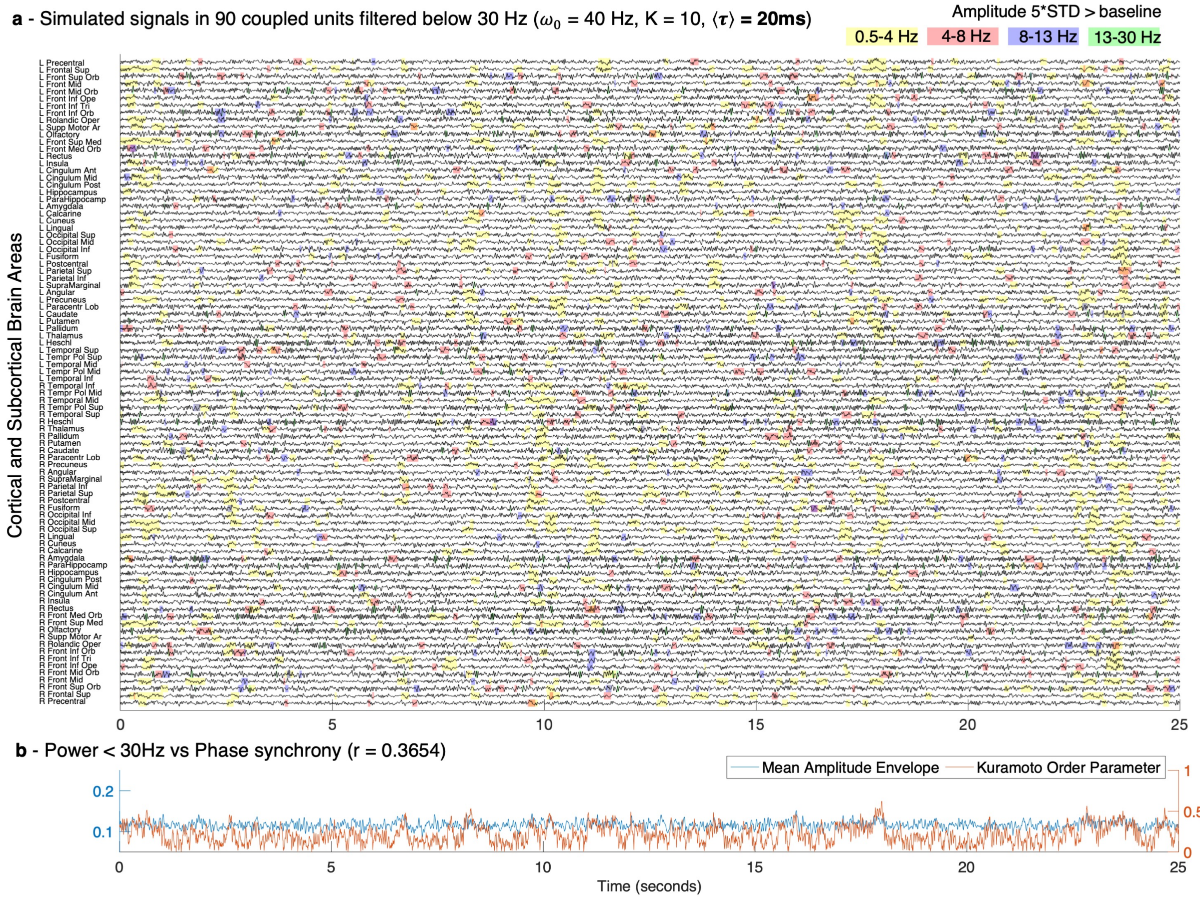


**Supplementary Figure 13. Detection of MOMs for long delays and intermediate coupling.** The simulated signals over 25 seconds in all 90 units, filtered below 30 Hz to concentrate on the sub-gamma oscillatory activity detected with MEG. Shades highlight the time points of increased power in the delta (yellow), theta (red), alpha (blue) and beta (green) frequency bands, which in this case do not emerge due to the weaknesses of the coupling strength. For each frequency band, the threshold was defined as 5 standard deviations (STD) from the amplitude in the same frequency bands of the simulated signals when no delays were considered (baseline simulations Supplementary Figure 11).

#### Supplementary Methods 1: Structural Connectivity

Long-range interactions between distant neuronal ensembles are mediated by a complex circuitry of long fibre bundles, the biological Connectome structure, that can be registered in-vivo and non-invasively using diffusion MRI [39-41]. We assume that the between-area connections designate the connections between those areas. All neuroimaging data were processed using two different parcellations: Shaefer[42] parcellation with 200 cortical brain regions and AAL parcellation, with 90 anatomically-segregated cortical and subcortical areas (excluding the cerebellum)[43]. We obtained the connectivity matrix $C_{N\times N}$ from two HCP dMRI dataset. The first dataset uses the highest quality multi-shell diffusion data acquired in sequence, taking 59 min from 985 HCP individuals (for HCP specifications, see their website http://www.humanconnectome.org/). The second dataset uses even better protocols, taking 89 min for each of 32 HCP participants at the Massachusetts General Hospital centre. Both dMRI datasets were preprocessed and made available as part of the freely available Lead-DBS software package (http://www.lead-dbs.org/).

Briefly, each entry $C_{np}$ in the connectivity matrix $C_{N\times N}$ is scaled in proportion to the number of fibres detected with tractography between each pair of brain areas $n$ and $p$, with $n, p \in N$. Importantly, the matrix $C$ is organized in a way that equivalent contralateral regions are arranged symmetrically, such that $C_{np} = C_{pn}$ (i.e., symmetric coupling).

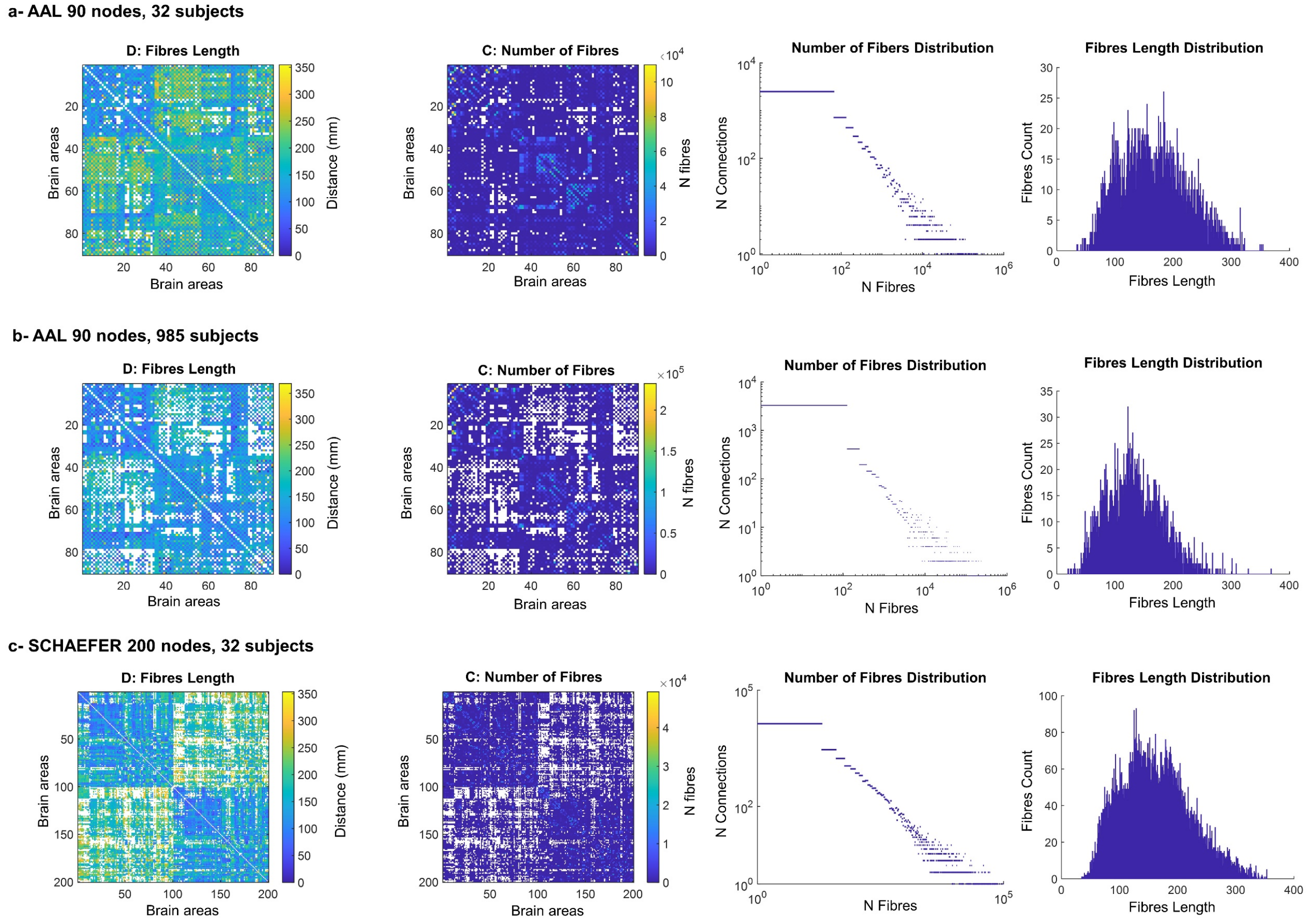


**Supplementary Figure 14. Anatomic Brain networks derived from Diffusion Tensor Imaging (DTI) from the Human Connectome Project public database. a-** AAL parcellation, 90 brain areas, averaged across 32 HCP healthy subjects. **b-** AAL parcellation, 90 brain areas, averaged across 985 HCP healthy subjects. **c-** Shaefer parcellation, 200 brain areas, averaged across 32 HCP healthy subjects. For each of the three profiles: (left) the matrix of fibre lengths D is calculated as the average length (in millimetres) of all fibres detected between each pair of regions using tractography, while the connectivity matrix C shows the average number of fibres detected between any pair of regions of interest demarcated according to the specific parcellation across healthy subjects; (right) distribution of coupling weights from C and distribution of fibre lengths from D, which is scaled by the mean delay to obtain the matrix of time delays.

#### *Validation of theoretical predictions across different connectomes*

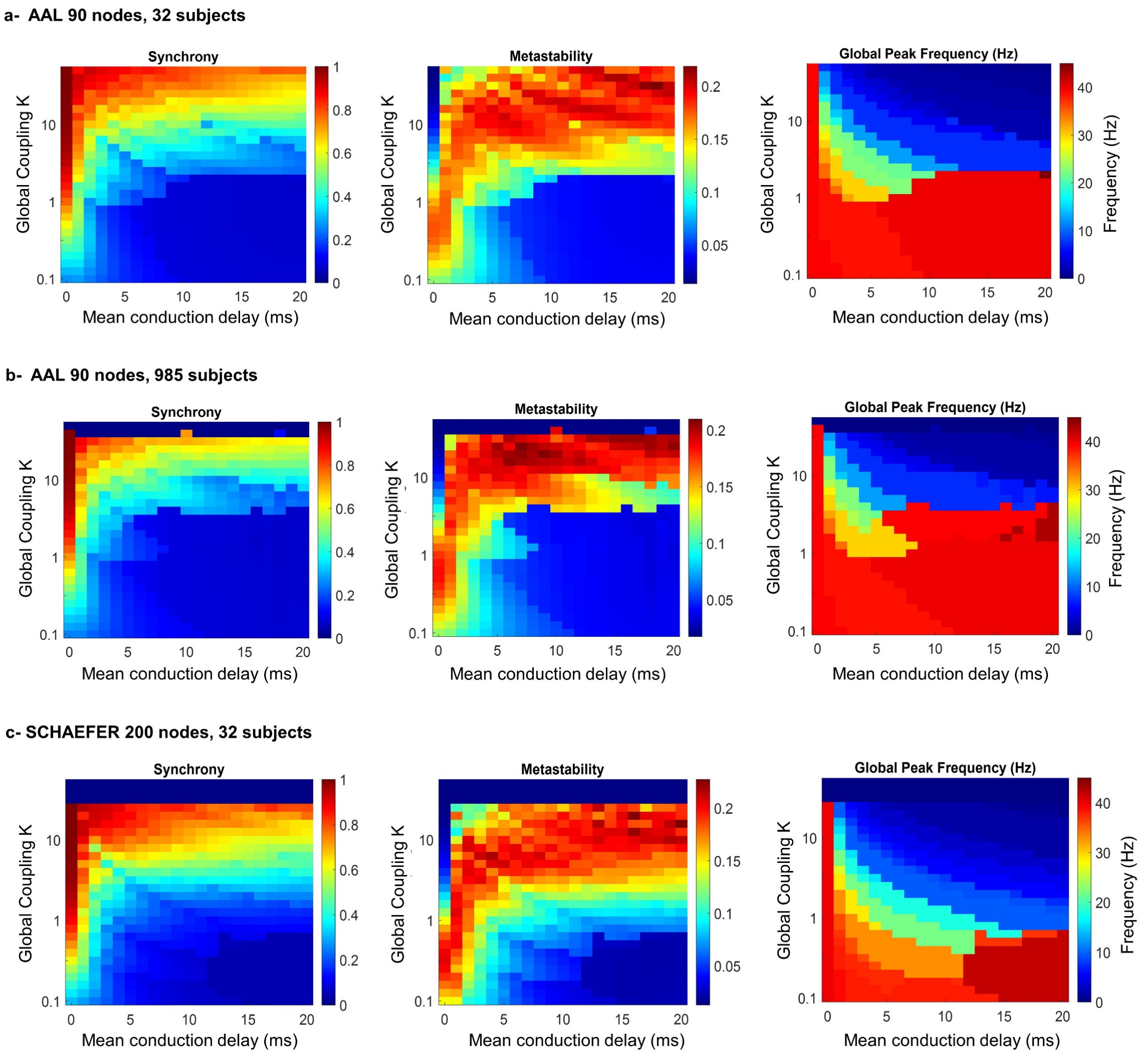


**Supplementary Figure 15. Theoretical predictions are consistent across different connectome profiles. Synchrony, metastability and global peak frequency are measured for three different connectome frameworks. a-** AAL parcellation, 90 brain areas, averaged across 32 HCP healthy subjects. **b-** AAL parcellation, 90 brain areas, averaged across 985 HCP healthy subjects. **c-** Shaefer parcellation, 200 brain areas, averaged across 32 HCP healthy subjects.

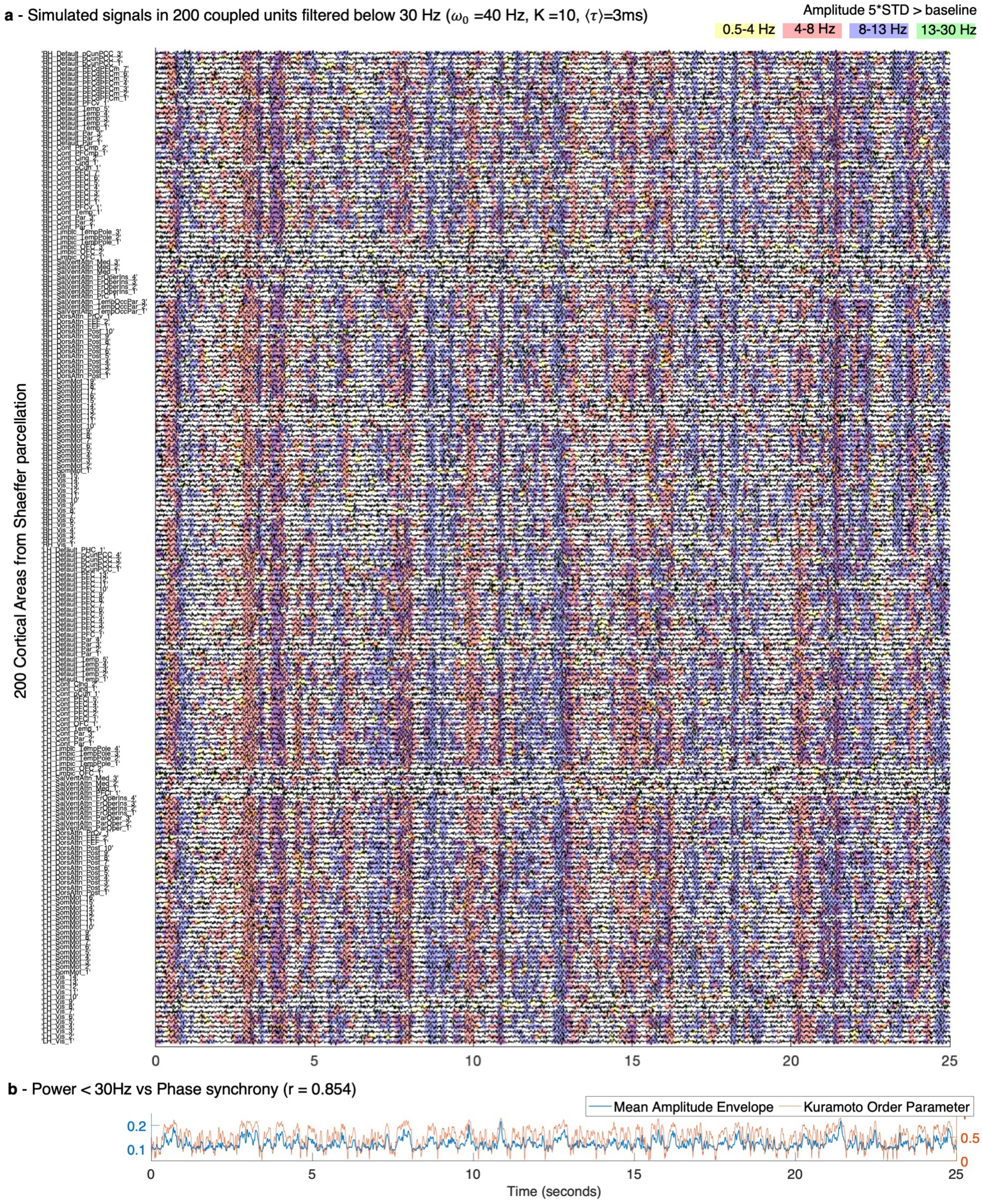


**Supplementary Figure 16. MOMs analysis in one optimal point across 200 brain cortical regions. a –** An example of the simulated signals in all 200 units plotted over 25 seconds, each representing a brain area from the Shaefer brain parcellation template, filtered below 30 Hz to highlight the sub-gamma oscillatory activity typically detected with MEG. Shades indicate the time points of increased power in the delta (yellow), theta (red), alpha (blue) and beta (green) frequency bands. For each frequency band, the threshold was defined as 5 standard deviations (STD) of the amplitude – in the same frequency bands – when no delays were considered. **b –**The mean amplitude envelope (blue) of the filtered signals shown in panel A correlates with r=0.854 with the phase synchronization evaluated by the Kuramoto Order Parameter (orange, right y-axis).

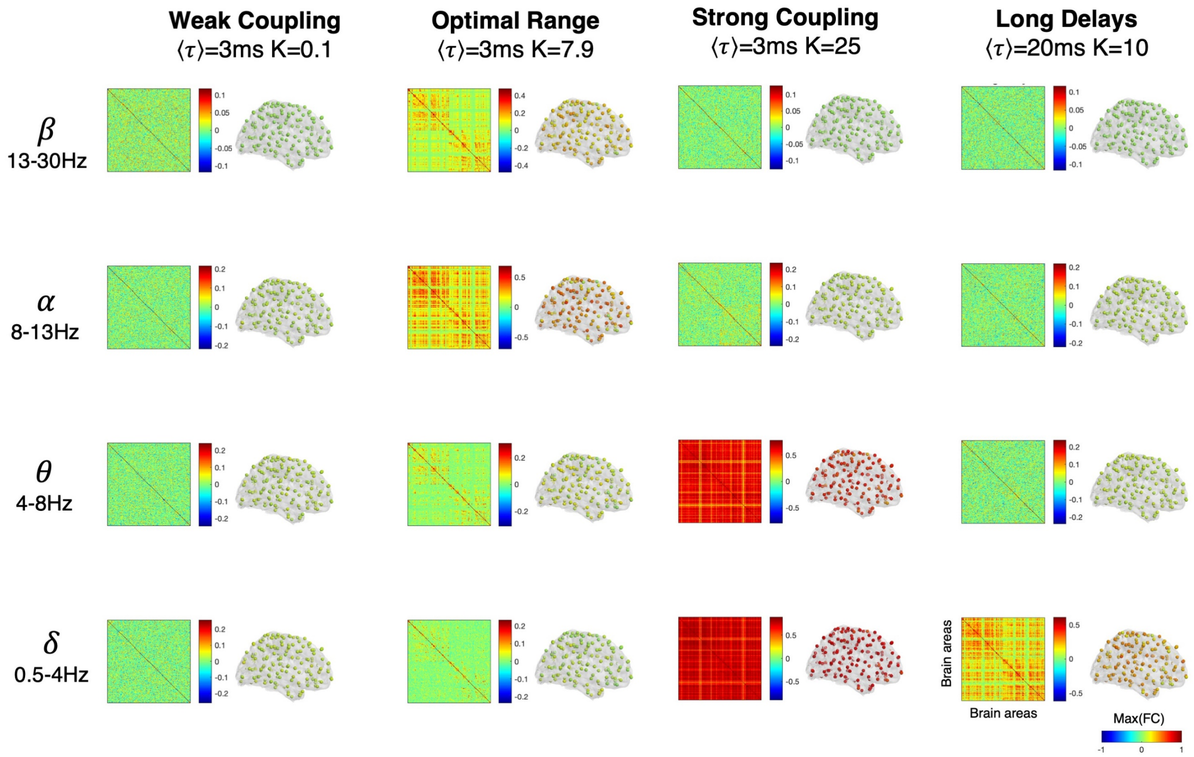


**Supplementary Figure 17. Repertoire of frequency-specific envelope functional connectivity patterns emerging from synchronization in the connectome using the Shaefer parcellation into 200 cortical-only brain areas.** For different sets of model parameters we report the frequency-specific envelope connectivity.

*Resolution of numerical integration*

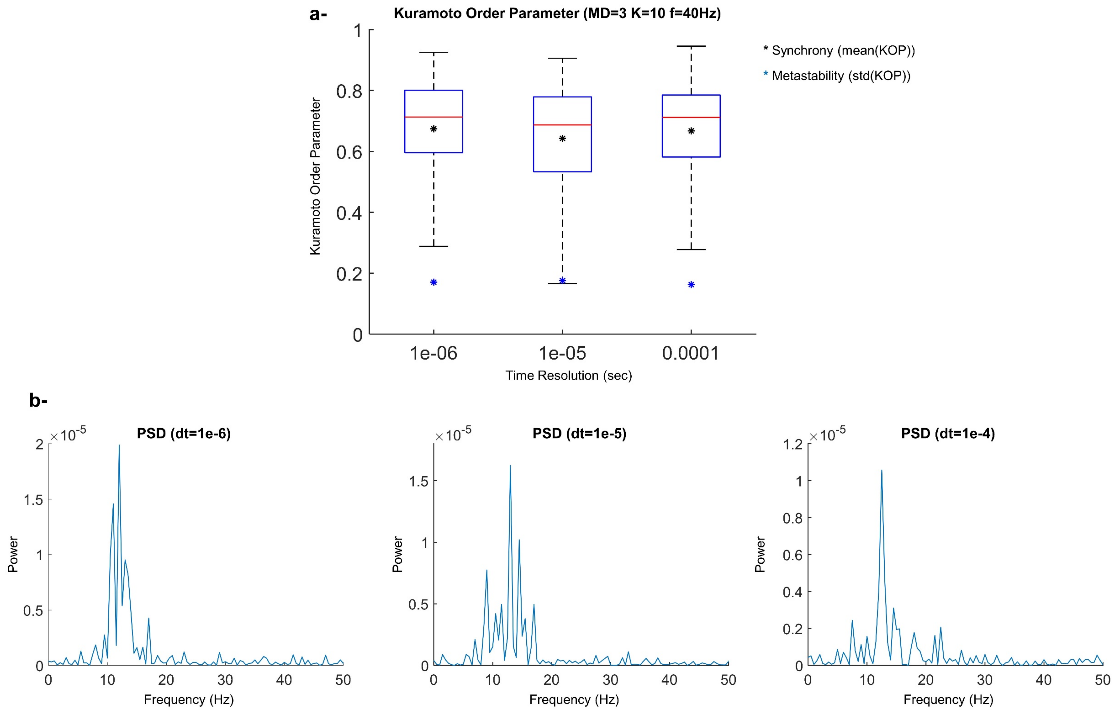


**Supplementary Figure 18. Effect of the integration time steps value on the model global parameters.** To evaluate how sufficiently small the integration steps must be for the model to capture the relevant global dynamical features, we perform simulations with integration time steps of dt=1e-6, 1e-5, 1e-4 in one of the optimal points (MD=3, K=10). For each simulation: **a-** Kuramoto Order Parameter (KOP) is measured across N=90 nodes, and synchrony and metastability is reported (black and blue asterisks, respectively). **b-** The global peak frequency averaged across nodes is computed. This analysis shows that a time resolution of dt=1e-4 would suffice for exploring the whole network parameter space, since the level of synchrony and metastability in the system (mean(KOP) and std(KOP), respectively) are not significantly impacted, and the global peak frequency is around the same value (here 11-13Hz).

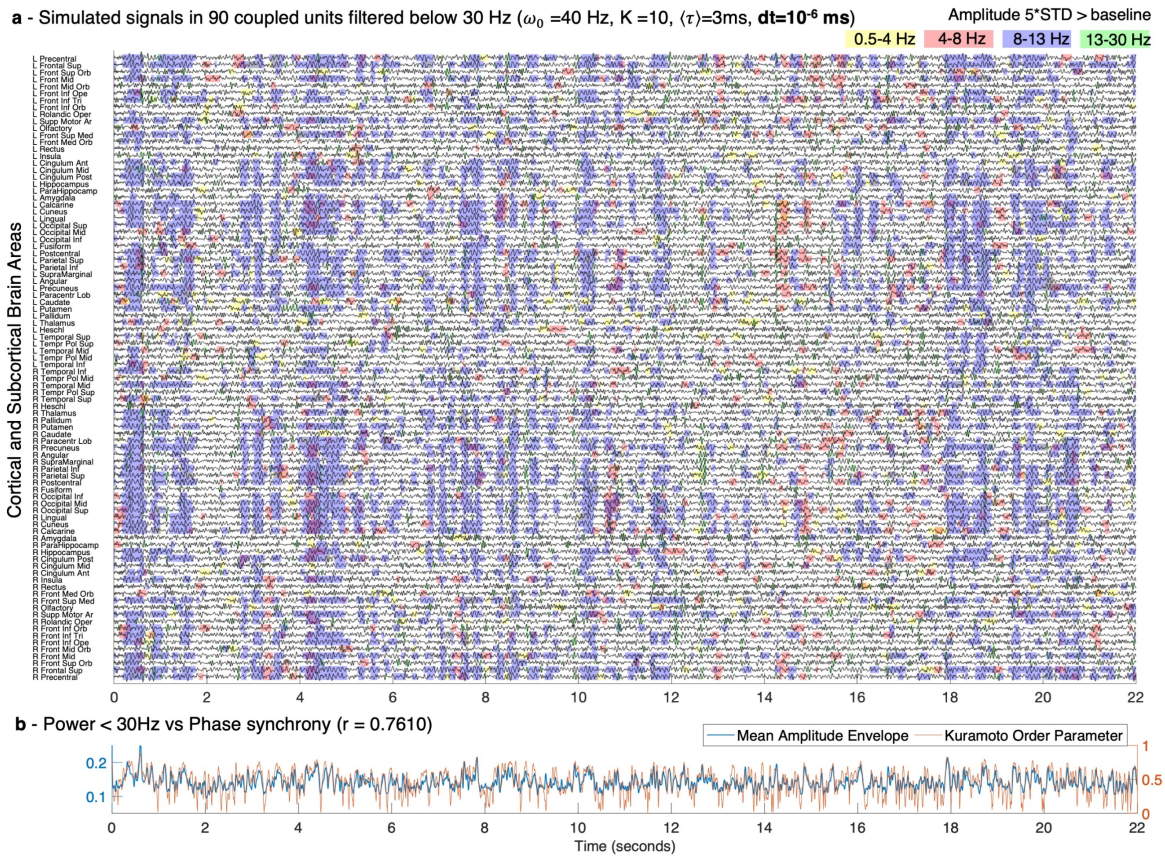


**Supplementary Figure 19. a – Metastable Oscillatory Modes (MOMs) detected for simulations with a reduced time step of numerical integration ( (i.e., dt=$10^{-6}$ instead of dt=$10^{-4}$).** Simulated signals in all 90 units for the same parameters used in Figure 4 of the manuscript but reducing the time step of numerical integration by two orders of magnitude, showing consistency of the results, with MOMs emerging mostly in the alpha frequency range. **b –** In line with the results reported for a larger time step, the mean amplitude envelope (blue) of the filtered signals shown in panel A correlates with r=0.7610 with the phase synchronization evaluated by the Kuramoto Order Parameter (orange, right y-axis).

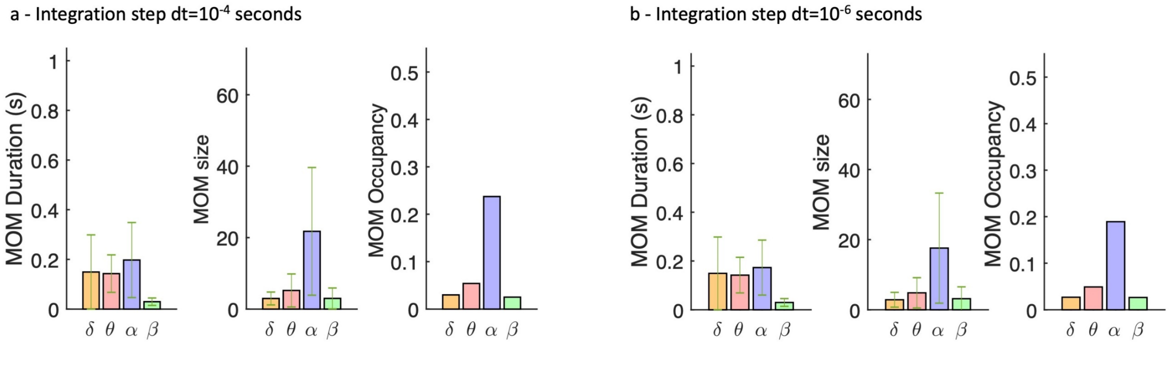


**Supplementary Figure 20. Characterization of MOMs for different integration time steps.** MOMs are characterized for two simulations performed with the same parameters but different time integration steps in terms of duration (i.e., consecutive time that the power remains above threshold), size (i.e., the number of units simultaneous displaying power above threshold) and occupancy (i.e., the proportion of time that the power is detected above threshold over the entire simulation), for each frequency band. The peak in alpha MOM size and occupancy is consistent in the two simulations, demonstrating that the results are not due to artifacts associated with the timestep for numerical integration.
